# Evolution of Modularity, Interactome and Functions of GIV/Girdin (CCDC88A) from Invertebrates to Vertebrates

**DOI:** 10.1101/2020.09.28.317172

**Authors:** Jason Ear, Amer Ali Abd El-Hafeez, Suchismita Roy, Tony Ngo, Navin Rajapakse, Julie Choi, Soni Khandelwal, Majid Ghassemian, Luke McCaffrey, Irina Kufareva, Debashis Sahoo, Pradipta Ghosh

## Abstract

PDZ domains are one of the most abundant protein domains in eukaryotes and frequently found on junction-localized scaffold proteins. Various signaling molecules bind to PDZ proteins *via* PDZ-binding motifs (PBM) and finetune cellular signaling. Here we describe the presence of a PBM on GIV/Girdin (CCDC88A) that is conserved throughout evolution, from invertebrates to vertebrates, and is generated as a *long* isoform-variant in humans, which we named *GIV-L*. Unlike GIV, which lacks PBM and is cytosolic, GIV-L localizes to the cell junctions, and has a unique PDZ-interactome, which impacts *GIV-L*’s ability to bind and activate trimeric G-protein, Gi through its *<u>g</u>*uanine-nucleotide *<u>e</u>*xchange *<u>m</u>*odulator (GEM) module; the GEM module is found exclusively in vertebrates. Thus, the two functional modules in GIV evolved sequentially: the ability to bind PDZ proteins via the PBM evolved earlier in invertebrates, whereas G-protein binding and activation may have evolved later only among vertebrates. Phenotypic studies in Caco-2 cells revealed that GIV and GIV-L may have antagonistic effects on cell growth, proliferation (cell cycle), and survival. Immunohistochemical analyses in human colon tissues showed that GIV expression increases with a concomitant decrease in GIV-L during cancer initiation. Taken together, these findings reveal how GIV/CCDC88A in humans displays evolutionary flexibility in modularity, which allows the resultant isoforms to play opposing roles either as a tumor suppressor (GIV-L) or as an oncogene (GIV).

## Introduction

Scaffolding proteins are important molecules that regulate the temporal, spatial and kinetic aspects of protein complex assembly (Scott and Pawson 2009, Rouaud, Sluysmans et al. 2020). Advantaged by their multi-modular makeup (Pawson and Nash 2003), they modulate the local concentrations of, proximity to, and subcellular dispositions and biochemical properties of the target proteins to impart plasticity within dynamic, spatially restricted intracellular signaling, earning them the reputation of ‘placemakers’ and ‘pacemakers’ of cell signaling (Pan, Sudol et al. 2012). Among the numerous modules that facilitate scaffolding, PDZ domains [Post synaptic density protein (PSD95), Drosophila disc large tumor suppressor (Dlg1), and Zonula occludens-1 protein (zo-1)] comprises one of the largest, and is frequently encountered in proteins at the cell junctions (Amacher, Brooks et al. 2020). PDZ binding motifs (PBMs) are short linear motifs commonly found on the C-terminus of proteins (although internal PBMs do exist) and mediate the PDZ•PBM interactions.

Members of the CCDC88 family of proteins are multi-modular molecular scaffolds that serve as signal transducers in eukaryotes (Enomoto, Murakami et al. 2005, Le-Niculescu, Niesman et al. 2005). In mammals, this family is comprised of three members: CCDC88A/GIV, CCDC88B/GIPIE, and CCDC88C/Daple; each member features a conserved HOOK-like domain and a coiled-coil domain on their N-terminal end. It is the C-termini of these that show more sequence divergence as well as a divergent modular makeup. Both GIV and Daple have a disordered C-terminal region which contain a G-protein exchange and modulator (GEM) motif which they use to bind and activate the heterotrimeric G-proteins of the Gi sub-family (Coleman, Marivin et al. 2016). GIPIE, on the other hand, has a shortened C-termini and lacks the GEM motif. Daple is unique from GIV in that it contains a PBM which allows it to bind to the regulator of Wnt signaling disheveled (Dvl) and a Frizzled binding module to bind Frizzled family of Wnt sensors (Oshita, Kishida et al. 2003, Aznar, Midde et al. 2015); in doing so, Daple links G-protein signaling to Wnt/Fzd pathways. (Oshita, Kishida et al. 2003, Enomoto, Murakami et al. 2005, Le-Niculescu, Niesman et al. 2005, Coleman, Marivin et al. 2016). The PBM module on Daple is also required for its localization at cell-cell junctions, *via* its ability to bind PDZ domain containing junctional proteins (PARD3, etc.) and such localization appears to be phosphomodulated (Marivin and Garcia-Marcos 2019, Ear, Saklecha et al. 2020).

Like Daple, GIV has also been observed at cell-cell junctions, and has been found to interact with junction-associated polarity proteins in mammalian cells (Houssin, Tepass et al. 2015, Sasaki, Kakuwa et al. 2015, Aznar, Patel et al. 2016, Siletti, Tarchini et al. 2017). How it may do so has remained unclear, especially because GIV in vertebrates does not appear to have the PBM module (that is seen in Daple). Interestingly, in invertebrates such as C. elegans and Drosophila (which lack Daple), GIV lacks its GEM motif, but contains a PBM which has been shown to regulate cilia function in C. elegans (Ha, Polyanovsky et al. 2015, Sasaki, Kakuwa et al. 2015, Nechipurenko, Olivier-Mason et al. 2016).

Here we report the discovery of a novel isoform of GIV in vertebrates (zebrafish and higher mammals), one which contains both, the G protein modulatory GEM motif as well as a conserved C-terminal PBM. This isoform not only offers insights into the evolution of the gene between invertebrates and vertebrates, but also sheds light on mechanisms that help enrich GIV onto cell-cell junctions. Finally, through meta-analysis of publicly available interaction data and through GIV biotin proximity labeling (BioID) we identified a GIV-PDZome interaction network. We propose that the compartmentalization of the two GIV isoforms may explain the dual role of GIV as both a tumor suppressor and an oncogenic driver, as previously reported in the literature (Ghosh, Beas et al. 2010).

## Results

### GIV has a PDZ-binding motif that is evolutionarily conserved among vertebrates and invertebrates

Prior characterization of GIV in invertebrate species such as C. elegans and drosophila described the presence of a PBM on the protein’s C-terminal end (Nechipurenko, Olivier-Mason et al. 2016). When we performed a BLAST alignment of the C. elegans and drosophila PBM (H_2_N-**EYGCV**-COOH) into the vertebrate database, we found that the PBM sequence aligned to several predicted GIV transcripts **(Figure 1A, Figure S1A and B)**. This intrigued us because of two reasons: First, this PBM sequence is highly conserved across all vertebrate and invertebrate species analyzed **(Figure S1B)**, suggesting a conserved evolutionary function. Second, despite such conservation, all studies on mammalian GIV until now used constructs lacking this module. It is also worth noting that GIV’s PBM sequence also resembles that of Daple’s **(Figure 3A)**, a gene that belongs to the same family as GIV, i.e., ccdc88 family (Oshita, Kishida et al. 2003, Nechipurenko, Olivier-Mason et al. 2016). This is particularly interesting because Daple, to the best of our knowledge, has not been found to exist in the genome of invertebrates **(Figure 1A and Figure S1A)**.

**Figure 1.**
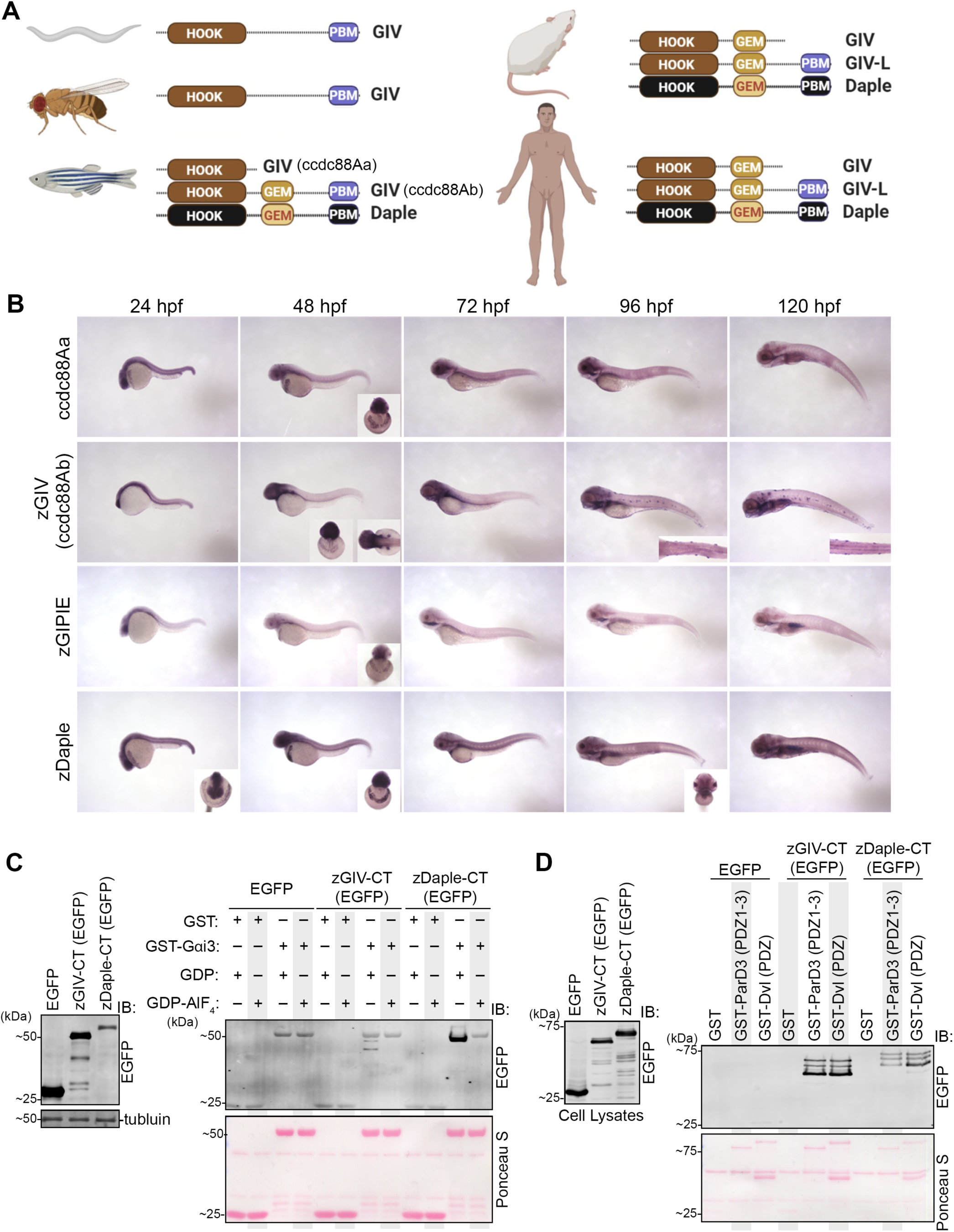
The C-terminus of GIV has an evolutionarily conserved functional PDZ-binding motif downstream of its G protein binding and/or modulatory domains. **A)** Schematic depicting the major modules and motifs within GIV and Daple across different species. HOOK, a highly conserved microtubule-binding NH2-domain; GEM, guanine nucleotide-exchange modulator; PBM, PDZ-binding motif; GIV-L, long isoform of GIV. **B)** Whole-mount RNA in situ hybridization of the CCDC88 gene family in developing zebrafish embryos across multiple time points. Inset shows anterior or dorsal view of select embryos. **C)** GST-pulldown assays were carried out using purified rat Gαi3 (loaded with GDP or GDP-AlF_4_^-^) and lysates of HEK293T cells exogenous expressing zebrafish GIV-CT or Daple-CT. Bound proteins were analyzed (right) and equal loading of lysates were confirmed (left) by immunoblotting (IB). **D)** GST-pulldown assays were carried out using purified GST-tagged PDZ domains of ParD3 and Dvl and lysates of HEK293T cells exogenous expressing zebrafish GIV-CT or Daple-CT and bound proteins were visualized as in C.

**Figure 2.**
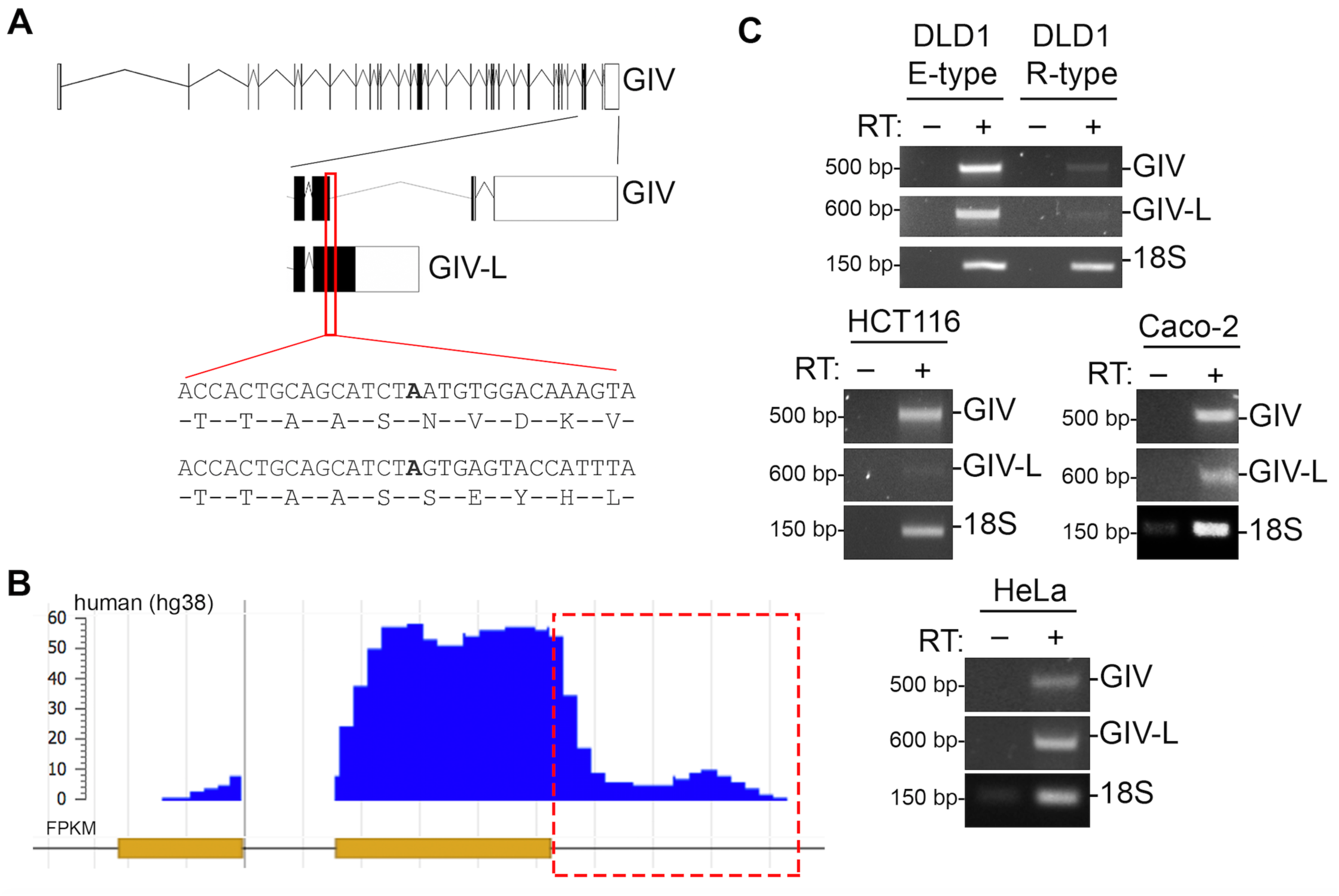
GIV-L, a human transcript for GIV, translates a variant protein that contains a PBM. **A)** Schematic of GIV and GIV-L transcript as annotated in ensembl. Indicated in red is region where transcript sequence of GIV (Exon 31) and GIV-L diverges. The panel below shows nucleotide sequence and translated sequence of indicated region. **B)** m6a-methylation sequence of GIV transcript as annotated in MeT-DB (http://compgenomics.utsa.edu/methylation/). Highlighted in red is corresponding region of GIV-L transcript (the intron immediately downstream of exon 31) indicating the methylated region. **C)** Reverse-transcription PCR of GIV and GIV-L transcript in multiple cell lines: DLD1 E-type and R-type (top), HCT116 (middle, left), Caco-2 (middle, right) and HeLa (bottom).

**Figure 3.**
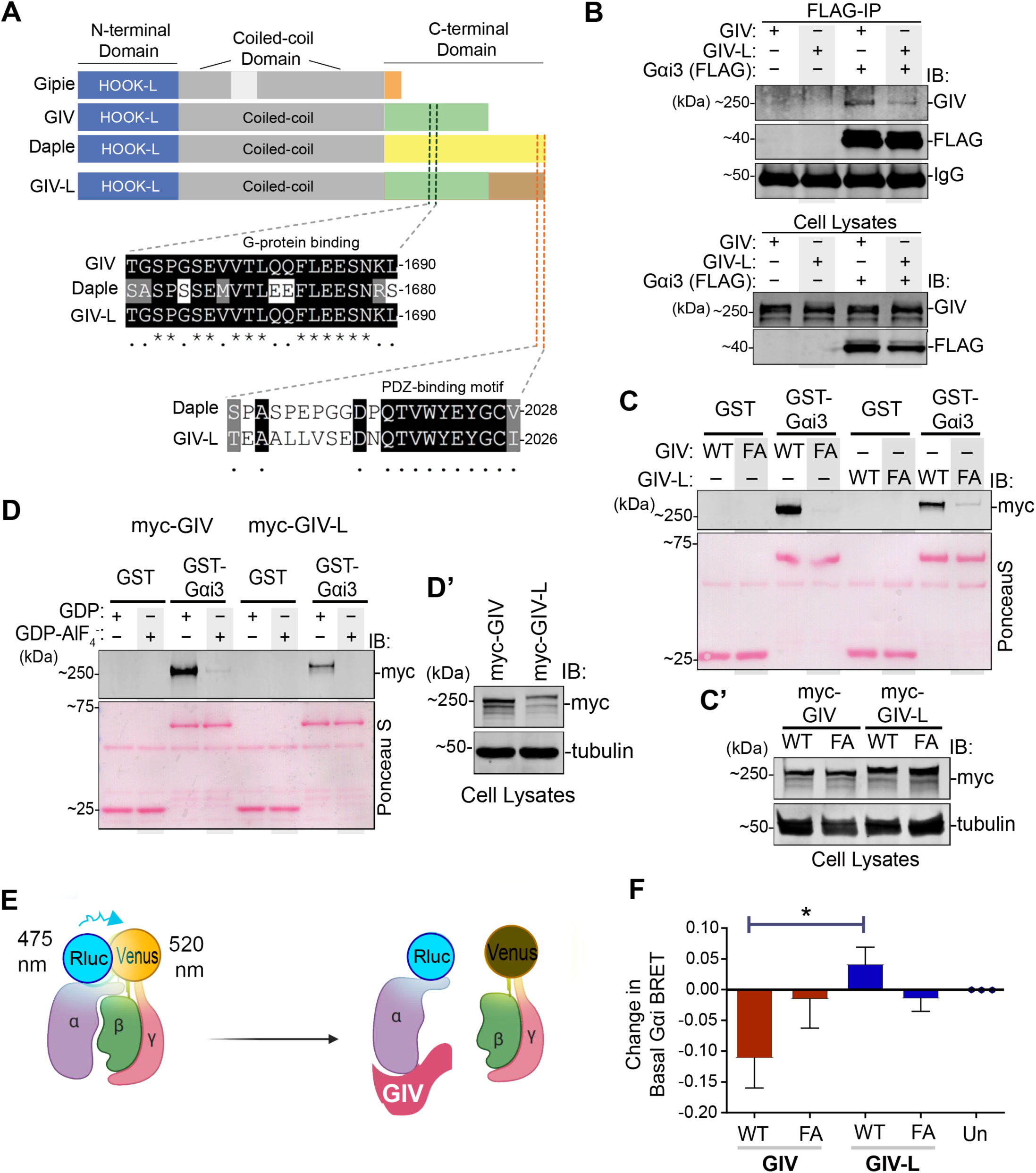
Both GIV and GIV-L use their GEM motifs to preferentially bind GDP-bound Gαi, but only GIV WT, and not other GIV variants, reduce basal Gai-RLuc2/mVenus-Gbg BRET in HEK293T cells. **A)** A schematic displaying the modular makeup of the CCDC88 family of proteins, from top to bottom-- CCDC88A/GIV, CCDC88B/GIPIE, and CCDC88C/Daple. **B)** Equal aliquots of lysates of HEK293T cells co-expressing FLAG-tagged Gαi3 and either myc-tagged GIV or GIV-L constructs were subjected to immunoprecipitation assays using an anti-FLAG antibody. Bound proteins and cell lysates were assessed for Gαi3 (FLAG) and GIV (myc) by immunoblotting (IB). **C)** GST-pulldown assays were carried out using purified GST-Gαi3 and lysates of HEK293T cells exogenous expressing myc-tagged wild-type (WT) or F1685A mutant (FA) of human GIV or GIV-L. Bound GIV was analyzed by immunoblotting (IB) with an anti-myc antibody. Panel C’ shows expression of proteins in the HEK293T cell lysates that were used as source of GIV in pulldown assays. **D)** GST-pulldown assays were carried out using purified GST-Gαi3, pre-loaded with GDP or GDP-AlF_4_^-^, and lysates of HEK293T cells exogenously expressing myc-tagged wild-type human GIV or GIV-L. Panel D’ shows expression of proteins in the HEK293T cell lysates that were used as source of GIV in pulldown assays. **E)** A schematic representation of the Gαi1(91)-RLuc2/mVenus-Gβγ BRET experiment. In the Gαiβγ heterotrimer, the proximity of RLuc2 (fused to Gαi1) to mVenus (fused to Gβγ) generates higher energy transfer (BRET); reduced BRET indicates the dissociation of Gαi1(91)-RLuc2 from mVenus-Gβγ. **F)** Change in basal Gαi1(91)-RLuc2/mVenus-Gβγ BRET in HEK293T cells transfected with the indicated GIV variants, compared to untransfected cells in the same experiment. The average BRET was calculated over 3 mins after adding the Rluc2 substrate, Coelenterazine-h, and the corresponding value from untransfected cells was subtracted. The experiment was performed in three independent biological replicates on different days, each containing three technical replicates. Error bars represent S.E.M. (n = 3 biological replicates). The graphs were plotted using GraphPad Prism 5 and statistical significance was calculated using one-way Anova with Tukey’s Multiple comparisons test.

To further characterize GIV’s PBM, we first analyzed the expression pattern of GIV (and the other ccdc88 family members) in zebrafish embryos. We chose zebrafish because: 1) it is a vertebrate animal in which all three members of the ccdc88 family exists-A-C; 2) its small size and rapid development allows for analysis of ccdc88 expression in the entire intact animal and across multiple timepoints, and 3) a systematic study on GIV in zebrafish (or for that matter, any of the ccdc88 family members) has not been done. We noted that the GIV gene is duplicated in zebrafish and is annotated as ccdc88Aa and ccdc88Ab **(Figure 1A)**; such duplication is a frequent event in teleost evolution (Volff 2005). Whole-mount RNA in situ hybridization on zebrafish embryos between 24- and 120-hours post fertilization (hpf) reveal a ubiquitous expression pattern for all ccdc88 family members at 24 hpf **(Figure 1B)**. As development progresses, expression is localized towards the anterior region of the embryo, on structures that appear to be hatching gland cells over the yolk sac at 48 hpf. Only ccc88Ab shows expression onto structures resembling lateral line hair cells at 96 and 120 hpf **(Figure 1B and Figure S1C)**. We selected ccdc88Ab (herein referred to as zGIV) for further studies due to its unique expression pattern and because it contains both the previously defined G protein regulatory GEM motif and the newly described conserved PBM sequence.

In order to confirm if the conserved GEM and PBM sequence on zGIV can indeed bind to the α-subunit of trimeric Gi-proteins and PDZ proteins, we overexpressed the C-terminus of zGIV tagged to EGFP in cells and subjected the cell lysates to interaction assays with purified GST-tagged Gαi3 **(Figure 1C)** or the PDZ domains of ParD3 and Dvl **(Figure 1D)**. The cell polarity regulator, ParD3, and the Wnt signaling regulator, Dvl, were chosen as the PDZ proteins because they have been identified as interactors of GIV in prior studies without a clear understanding of the mode of these interactions (Ohara, Enomoto et al. 2012, Sasaki, Kakuwa et al. 2015, Boldt, van Reeuwijk et al. 2016). In addition, these two PDZ proteins have been demonstrated to bind to Daple’s PBM (Oshita, Kishida et al. 2003, Aznar, Midde et al. 2015, Siletti, Tarchini et al. 2017, Ear, Saklecha et al. 2020). Finally, because of the high sequence similarity between GIV-L’s and Daple’s PBM, we rationalized that GIV-L’s PBM can also bind to ParD3 and Dvl. As a positive control, we tested the C-terminus of zebrafish Daple (zDaple) alongside zGIV in the same assays. Both proteins bound Gαi3 and PDZ proteins (**Figure 1C-D**). Furthermore, consistent with what is expected for GEMs, both zGIV and zDaple bound G-proteins in a nucleotide-dependent manner, in that, both proteins preferentially bound the inactive conformation of G proteins in presence of GDP, but not its active conformation mimicked by pre-loading the G-protein with GDP and aluminum fluoride, AlF_4_^-^ **(Figure 1C)**. Overall, these findings indicate that the PBM on zGIV is functional.

Taken together, we conclude that while GIV’s PBM module is functionally conserved in both vertebrates and invertebrates, the absence of a GEM motif in invertebrates suggests sequential evolution of the two functional modules: the ability to bind PDZ proteins via the PBM evolved earlier whereas G-protein binding and activation may have evolved later only among vertebrates.

### Human GIV has a long transcript isoform (GIV-L) coding for a C-terminal PBM

Several transcriptional isoforms of human GIV is predicted to have a PBM in its translated product **(Figure 1A, Figure S1A and B)** (XM_005264418.5, XM_017004476.2, XM_017004477.2). A closer analysis of the GIV transcript sequence revealed that the intron immediately downstream of exon 31 codes for an extended translated product **(Figure 2A)**. On the 3’ end of exon 31 of GIV, a splice event occurs which leads to the processing of the nascent transcript into GIV as describe in the current literature (NM_001135597.2) (Enomoto, Murakami et al. 2005, Le-Niculescu, Niesman et al. 2005). A new isoform of GIV lacks the splice event and, thus, leads to a processed transcript which translates to a larger GIV (by approximately 150 amino acids) protein that contains a PBM, inspiring the nomenclature, GIV-L.

Next we probed what might be triggers for alternative splicing of GIV into two isoforms. RNA methylation is a frequent event in eukaryotic cells and can affect RNA stability and splicing (Zhu, Zhu et al. 2018, Zaccara, Ries et al. 2019). Furthermore, N^6^-Methyladenosine (m^6^A) modification tend to occur on the last exon of a gene (Ke, Alemu et al. 2015). Interestingly, when we analyzed the methylation of GIV in a methyltranscriptome database (MeT-DB V2.0) (Liu, Flores et al. 2015, Liu, Wang et al. 2018) we observed that the same intronic region downstream of exon 31 is subjected to m^6^A modification **(Figure 2B)**. These analyses predict that mammalian cells have a protein coding transcript for an isoform of GIV which contains a PBM.

To validate the prediction, we designed unique primers to GIV and GIV-L (see *Methods* for sequence). Among the cell lines tested, we observed that the DLD1 E-type, Caco-2, and HeLa cells contain appreciable levels of GIV-L, whereas DLD1 R-type and HCT116 cells do not **(Figure 2C)**. It is noteworthy that among the colorectal cancer cell lines tested, DLD1 E-type and Caco-2 cells, and to some extent HeLa cells form cell-cell junctions whereas DLD1 R-type and HCT116 do not (Vermeulen, Bruyneel et al. 1995, Ilyas, Tomlinson et al. 1997), suggesting that GIV-L may be expressed and functional in cells with junctions.

### Both GIV and GIV-L can bind Gαi but with different affinities and/or abilities to dissociate trimers in cells

GIV and GIV-L have an identical GEM motif, however, only GIV-L contains a PBM **(Figure 3A)**; we asked if and how the presence of the PBM module impacts G protein binding. We performed a series of biochemical assays that have been used previously to rigorously validate GIV’s ability to bind and activate G-proteins (Garcia-Marcos, Ghosh et al. 2010). Coimmunoprecipitation experiments confirmed that while both GIV and GIV-L bind G-protein, Gαi3 **(Figure 3B)**, binding is weaker in the case of GIV-L. GST pulldown assays using GST-tagged Gαi3 and cell lysates as source of GIV/GIV-L further confirmed that while both GIV isoforms bind the G protein, and both isoforms require a functional GEM motif to do so (as determined by the loss of binding with a well-characterized GEM-deficient F1685A mutant), there is less binding in the case of GIV-L **(Figure 3C** and **C’)**. Finally, we sought to determine if GIV-L, similar to GIV, bind to G-proteins in a GDP-dependent manner. When GST-tagged Gαi3 was purified and loaded with GDP or GDP-AlF_4_^-^, we observed that GIV-L specifically binds to Gαi3 in the GDP-bound state **(Figure 3D** and **D’)**. These biochemical studies demonstrate the conserved ability of GIV-L, similar to GIV, to bind to G-proteins *in vitro*.

Solved structures of GIV-bound Gαi have confirmed that GIV engages the SwII region of Gαi and shares binding determinants with Gβγ, indicating (de Opakua, Parag-Sharma et al. 2017, Kalogriopoulos, Rees et al. 2019). These studies provided a structural basis for the experimentally confirmed ability of GIV to displace Gβγ from Gαi in cells, where it dissociates the Gi heterotrimer and triggers Gαi and Gβγ signaling (Midde, Aznar et al. 2015). To determine if binding of GIV-L to Gαi that we observe in vitro actually translates into G protein signaling in cells, we utilized a bioluminescence resonance energy transfer (BRET) assay measuring the association between luciferase-tagged Gαi and mVenus-tagged Gβγ **(Figure 3E)** (Breton, Sauvageau et al. 2010, Brown, Lambert et al. 2016). In this assay, a loss of BRET signal indicates Gβγ dissociation from the G-alpha subunit and, therefore, activation of the G-protein. In HEK293T cells ectopically expressing GIV (wt or F1685A) or GIV-L (wt or F1685A), we see, as expected, a decrease in the BRET ratio only in the presence of GIV (wt) but not GIV (FA). Surprisingly, neither GIV-L wt, nor GIV-L-FA constructs showed a significant change in BRET **(Figure 3F)**. These findings suggested that despite the fact that both GIV and GIV-L have a functional GEM motif which can bind Gαi in vitro, only GIV, but not GIV-L may trigger G protein signaling in cells under the conditions tested.

### Multiple PDZ proteins bind the PBM on GIV-L and modulate Gα-protein binding

A notable feature of the C-terminal PBM on GIV-L is its highly conserved sequence, which, like that in Daple **(Figure 3A)**, includes two tyrosine residues. Those sites in Daple serve as substrates for multiple tyrosine kinases and their phosphorylation has been shown to modulate PDZ•PBM interactions with ParD3 and Dvl **(Figure 4A)** (Aznar, Midde et al. 2015, Ear, Saklecha et al. 2020). In coimmunoprecipitation experiments between GIV-L and ParD3, we observed a specific interaction between GIV-L and the third PDZ domain on ParD3 **(Figure 4B)**, mirroring exactly what was shown previously for Daple (Ear, Saklecha et al. 2020). This ability to bind multiple PDZs (apparent promiscuity), and yet, preferentially doing so *via* one but not the other PDZ module on the same protein (i.e., specificity) highlights a key property of the conserved PBM binding modality shared by GIV-L and Daple. As with Daple, Dvl co-immunoprecipitated exclusively with GIV-L, but not with GIV or a deletion mutant of GIV-L lacking the C-terminal PBM (GIV-L ΔPBM) **(Figure 4C)**. These findings show the similarities between GIV-L’s and Daple’s PBM and demonstrate that this module is necessary for the GIV-L•PDZ interactions.

**Figure 4.**
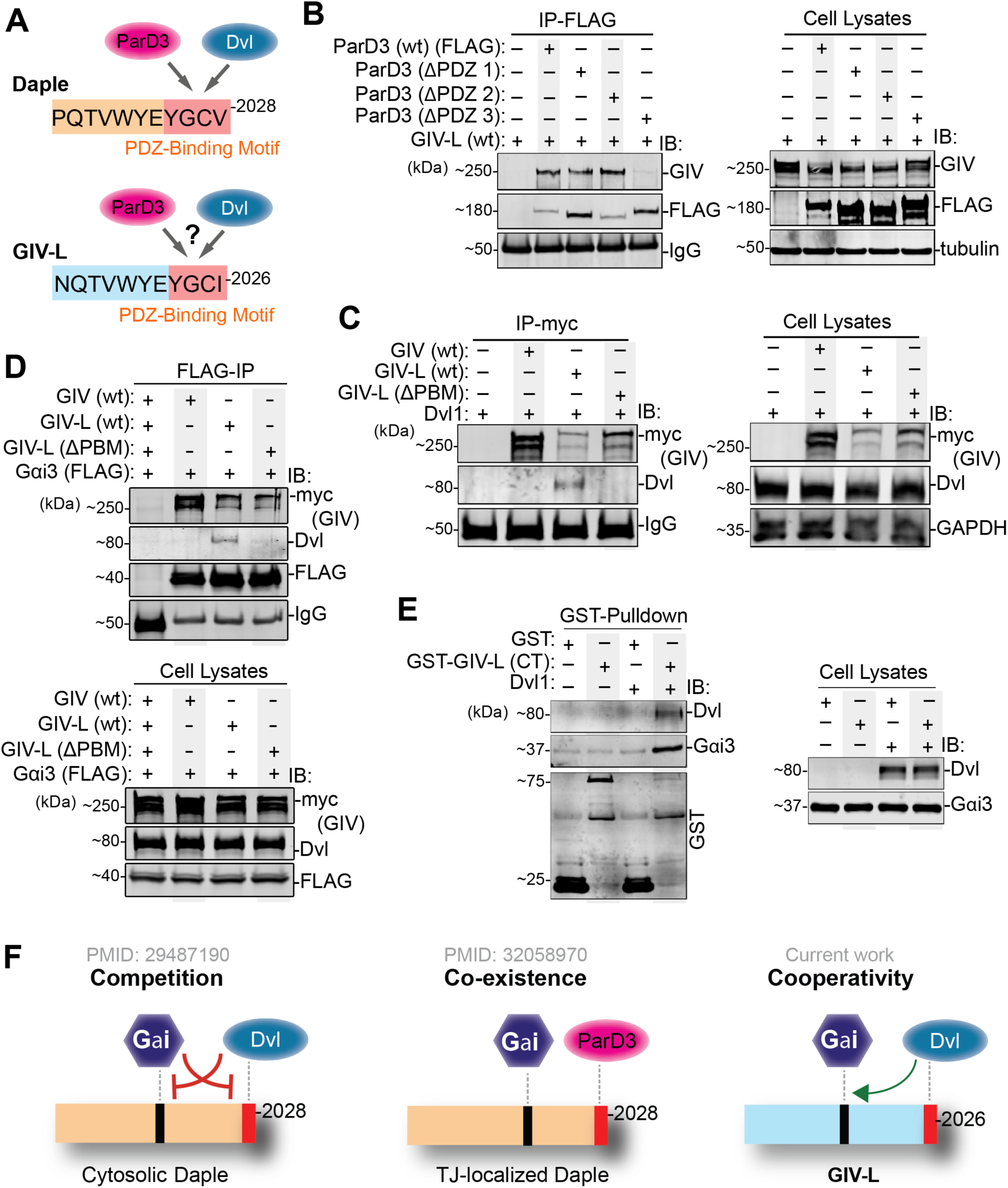
The PBM motif in GIV-L binds to multiple PDZ proteins and enhances G-protein binding. **A)** Schematic depicts the similarities between the sequences of the C-terminal PBMs (highlighted in red) of Daple and GIV-L and their respective immediate N-terminal flanking regions. While Daple’s PBM is known to bind ParD3 and Dvl, whether GIV-L can bind is tested here. **B)** Equal aliquots of lysates of HEK293T cells co-expressing various FLAG-tagged ParD3 constructs and GIV-L (wt) were subjected to immunoprecipitation assays using an anti-FLAG antibody. Bound proteins (left) and cell lysates (right) were assessed for ParD3 (FLAG) and GIV (myc) by immunoblotting (IB). **C)** Equal aliquots of lysates of HEK293T cells co-expressing untagged Dvl1 and either myc-tagged GIV (wt) or GIV-L (wt or ΔPBM) were subjected to immunoprecipitation assays using an anti-myc antibody. Bound proteins (left) and cell lysates (right) were assessed for Dvl and GIV (myc) by immunoblotting (IB). **D)** Equal aliquots of lysates of HEK293T cells co-expressing FLAG-tagged Gαi3 and either myc-tagged GIV (wt) or GIV-L (wt or ΔPBM) were subjected to immunoprecipitation assays using an anti-FLAG antibody. Bound proteins (top) and cell lysates (bottom) were assessed for Gαi3 (FLAG), GIV (myc), and endogenous Dvl by immunoblotting (IB). **E)** Equal aliquots of lysates of HEK293T cells co-expressing untagged Dvl and either GST or GST-tagged GIV-L (CT) was incubated with glutathione agarose beads. Bound proteins (left) and cell lysates (right) were assessed for Gαi3, Dvl, or GST by immunoblotting (IB). **F)** Schematic summarizing the differential impacts of binding of PDZ proteins to the C-terminal PBM motifs in Daple (left, middle) and GIV-L (right) on their ability to bind Gαi protein.

Prior work showed that binding of some PDZ proteins to Daple’s PBM modulates the ability of the latter to bind G proteins **(Figure 4F)**-- while Dvl competes with Gαi3 for binding Daple, ParD3 does not; the latter is capable of co-existing in ternary ParD3•Daple•Gαi complexes (Aznar, Ear et al. 2018, Ear, Saklecha et al. 2020). In the case of GIV-L, coimmunoprecipitation assays from cell lysates overexpressing Gαi3 and either GIV or GIV-L (wt or ΔPBM) showed that Dvl can also precipitate with the Gαi3•GIV complex **(Figure 4D)**. Furthermore, Dvl was not observed to immunoprecipitate with either GIV or the PBM-deficient mutant GIV-L (ΔPBM), indicating that Gαi3 interacts with Dvl only in the presence of GIV-L with an intact PBM. Interestingly, we also observed enhanced binding of GIV-L to Gαi3 in the presence of Dvl **(Figure 4E)**. Because such augmentation was not observed with the F1685A GIV-L mutant (cannot bind G protein; **Figure S2A**) our findings highlight the modular nature of this protein and, in this case, cooperativity between them (**Figure 4F**), i.e., binding of PDZ proteins to the PBM module augments binding of the GEM module to G proteins. Findings paint a picture in which PDZ proteins, by virtue of their ability to bind PBMs on GIV-L and Daple finetune the G protein regulatory function of the latter. Because the PBMs in both proteins are located ∼400 aa downstream of their respective GEM modules, in a stretch that has been deemed to be intrinsically disordered without any semblance to any folded module (Coleman, Marivin et al. 2016) it is likely that intermodular phenomenon of competition or cooperativity are mediated via allosteric mechanisms such as binding-induced conformational changes. This context-dependent exposure of the GEM module in GIV-L may, in part, be responsible for the observed lack of change in BRET between Gai-RLuc and mVenus-Gbg in GIV-L-expressing cells (**Figure 3F**).

### The PBM on GIV-L is required for localization at cell-cell junctions

PDZ proteins are highly enriched in cell junctions where they can serve as docking stations for proteins with PBMs (Fanning and Anderson 1999, Subbaiah, Kranjec et al. 2011, Manjunath, Ramanujam et al. 2018). GIV has not only been implicated in regulating cell-cell junction, it has also been observed to localize to cell junctions in vivo (Siletti, Tarchini et al. 2017). While several works have established the importance of Daple’s PBM in localizing Daple to cell junctions (Marivin and Garcia-Marcos 2019, Ear, Saklecha et al. 2020), how GIV localizes to cell junctions remains elusive. The first clue that GIV-L may be a junction-localized protein comes from cell fractionation studies on DLD1 E-type cell lysates **(Figure 5A)**. Immunoblotting for GIV in these cells revealed the presence of two bands in the post nuclear supernatant (PNS). Both S100 (cytosol) and P100 (membrane) pools derived from the PNS fraction also contain the observed two bands; however, when the P100 fraction was further separated into detergent soluble and detergent insoluble fractions, the lower of the two bands was specific to the detergent soluble fraction while the higher of the two bands was specific to the detergent insoluble fraction. The detergent insoluble fraction is known to be enriched in many cell junction proteins because they are typically highly resistant to detergent solubilization (Wong 1997, Tang and Brieher 2013, Kannan and Tang 2015). The presence of the higher GIV species in the detergent-insoluble pool suggest that this species may be GIV-L. Next, we ectopically expressed GIV or GIV-L (wt or ΔPBM) into HEK993T cells and performed similar fractionation studies **(Figure 5B)**. When crude membrane (P100) extract was separated into detergent soluble and detergent insoluble fractions, we see that GIV equally distributes between the two pools, while GIV-L (wt) has a greater enrichment into the detergent insoluble pool **(Figure 5B)**. Such an enrichment was diminished when the PBM was truncated. Complementing these fractionation studies, when GIV or GIV-L (wt or ΔPBM) was overexpressed onto DLD1 E-type cells (with endogenous GIV depleted using CRISPR/Cas9) we see that only GIV-L (wt) possessed the ability to localize onto cell-cell junctions, whereas GIV and the PBM-deficient mutant GIV-L (ΔPBM) remained cytosolic **(Figure 5C)**.

**Figure 5.**
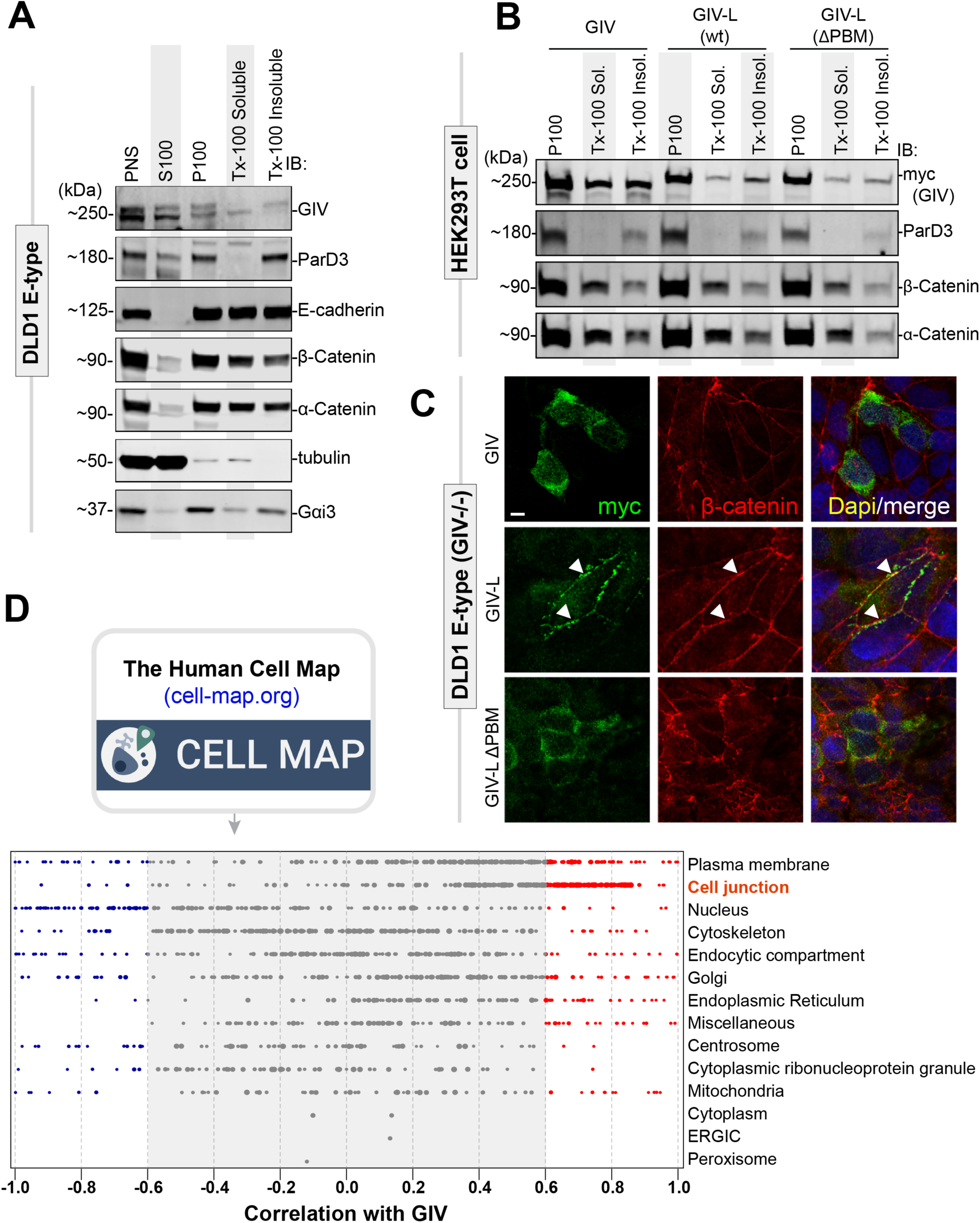
GIV-L, but not GIV, localizes at cell-cell junctions and its PBM is required for such localization. **A)** DLD1 E-type cells (parental or GIV-knockout) were fractionated into post nuclear supernatant (PNS), cytosolic (S100), crude membrane (P100), membrane detergent soluble (Tx-100 Soluble), and membrane detergent insoluble (Tx-100 Insoluble) pools. Equal proportions of each fraction was assessed for GIV by immunoblotting (IB). Equal loading and reasonable purity (lack of significant cross-contamination) of fractions was confirmed by immunoblotting for ParD3, E-cadherin, β-Catenin, α-Catenin, tubulin, and Gαi3. **B)** Cell fractionation studies were carried out as in **A** on HEK293T cells exogenously expressing myc-tagged GIV, GIV-L (wt), or mutant GIV-L (ΔPBM). Equal proportion of each fraction was assessed for GIV and other loading and/or fractionation controls (as above) by immunoblotting (IB). **C)** DLD1 E-type cells were transfected with myc-tagged GIV, GIV-L (wt), or mutant GIV-L (ΔPBM), methanol fixed, and stained with anti-myc (green) or anti-β-Catenin antibody. Arrowheads indicates cell-cell contact sites. Scale bar, 7.5 μm. **D)** The publicly available BioID-based proximity map of HEK293T cells annotated in the Human Cell Map (HCM), was queried for GIV (without distinguishing GIV and GIV-L) and other preys that co-traffic and localize and functionally associate with GIV (i.e., prey-prey correlations) for various organelle-specific baits. The interactome is enriched for cell junction-localized proteins.

Human Cell Map (HCM) is an extensive BioID-based proximity map of HEK293 cells using 192 compartment-specific baits (Go, Knight et al. 2019). Along with the proximity of preys to these compartment baits, prey co-trafficking, colocalization, and functional associations can be deduced from similarities in prey labeling by the array of baits, i.e. from prey-prey correlations (Go, Knight et al. 2019). The in-depth analysis of the HCM data indicated that GIV is highly correlated with proteins on cell junctions **(Figure 5D)** and, to a lesser degree, on plasms membrane. Taken together, these observations suggest that like Daple, GIV-L localizes to cell-cell junctions, and that such localization is enabled by its C-terminal PBM.

### BioID-proximity labeling identifies the PDZ-interactome (“PDZome”) of GIV-L

Because the modular composition of large scaffold proteins dictates the protein’s interactome, which in turn regulates its localization and functions, we next asked how the GIV interactome changed due to the additional PBM module in GIV-L. To this end, we carried out BioID proximity labeling coupled with mass spectrometry (MS) to identify GIV/GIV-L-interacting proteins **(Figure 6A)**. We validated our BirA-tagged GIV constructs using two approaches: first, we confirmed that these constructs can successfully biotinylate proteins *in situ* by incubating cell lysates with streptavidin beads followed blotting using fluorescent conjugated streptavidin **(Figure 6A)** and, second, we confirmed that the tagged constructs show the expected subcellular localizations **(Figure 6B)**. In agreement with what was observed in DLD1 E-type cells, GIV-L was found both in cytosol and at cell-cell contact sites, whereas GIV was largely cytosolic in localization.

**Figure 6.**
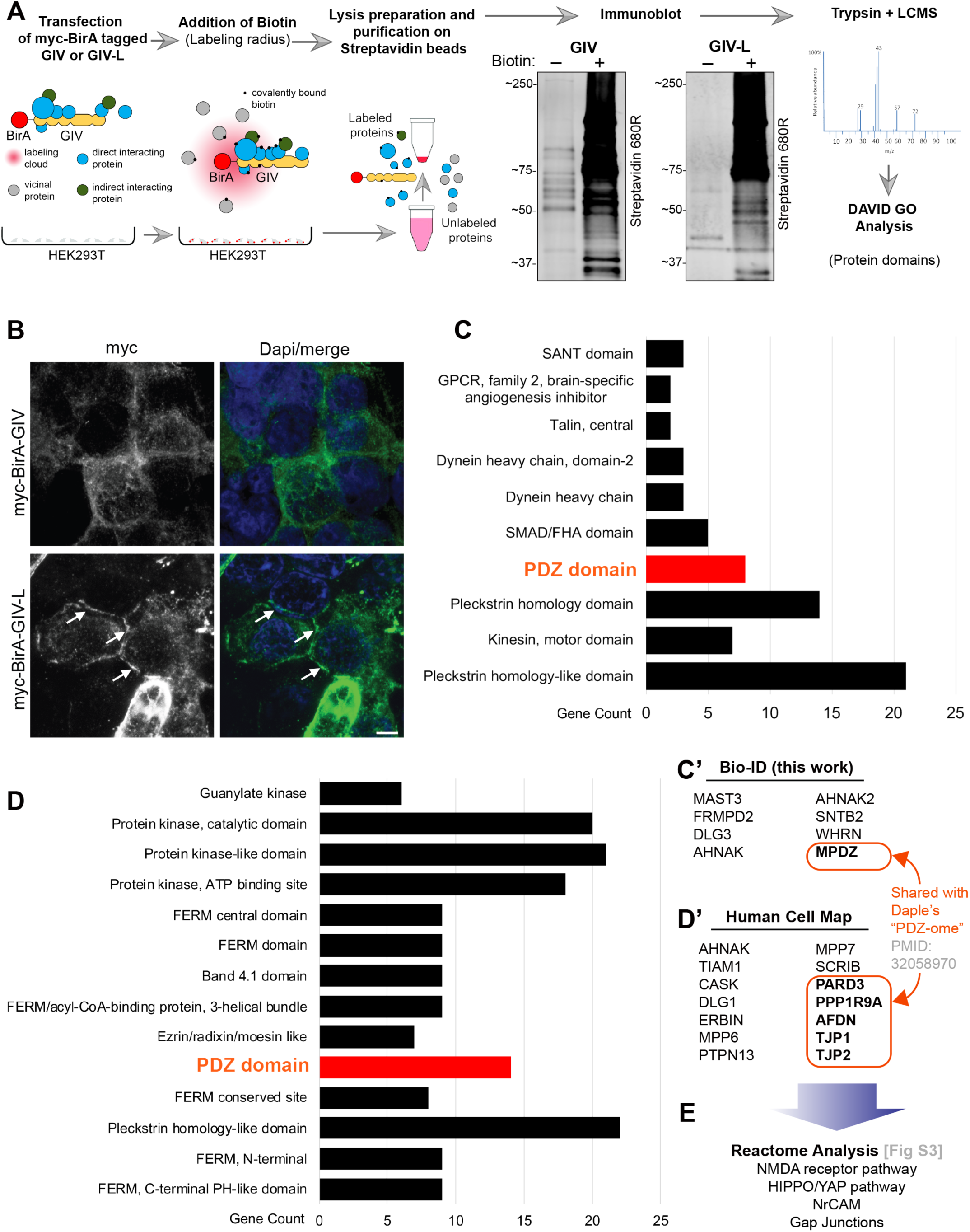
A protein-protein interaction (BioID) screen identifies the PDZ-interactome of GIV-L. **A)** Schematic depicting the key steps in biotin proximity labeling (BioID) studies used to identify the GIV and GIV-L interactomes in HEK293T cells. HEK293T cells were transiently transfected with myc-BirA tagged GIV or GIV-L construct and then treated with free biotin. Equal aliquots of cell lysates were incubated with streptavidin magnetic beads and proteins were eluted by boiling in the presence of excess free biotin. Eluted proteins were analyzed by SDS-PAGE and blotted with AlexaFluor-680 conjugated streptavidin to confirm successful proximity labeling. **B)** HEK293T cells exogenously expressing myc-BirA-tagged GIV or GIV-L were fixed with methanol prior to staining using anti-myc antibody. Arrows indicate localization onto points of cell-cell contact. Scale bar, 5 μm. **C)** Bar graph summarizing the GIV-L-interacting proteins (by mass spectrometry, grouped by protein domain using DAVID GO analysis. Top domains categories are shown. C’) List of PDZ domain proteins identified. **D)** Bar graph summarizing GIV’s interactome as annotated in the Human Cell map database, also grouped by protein domain using DAVID GO analysis. Top domains categories are shown. Panel D’ lists the PDZ-domain containing proteins reported in the Human Cell Map database.

Mass spectrometry identification and gene ontology analysis (via protein domain) of the biotinylated proteins in HEK293T revealed PDZ proteins proximity labeled with GIV-L, but not GIV **(Figure 6C and C’)**. Annotation and analysis of HCM data using our same gene ontology analysis revealed that there is indeed a unique set of “PDZome” that interacts with GIV-L **(Figure 6D and D’)**. Furthermore, Reactome pathway analysis performed on GIV-L’s PDZome identified overrepresentation of NMDA and HIPPO pathways—two pathways that are closes associated with junctional sensing (Wang, Denisova et al. 2010, Karaman and Halder 2018, Rausch and Hansen 2020).

### GIV is required for contact-dependent growth inhibition, cell-cycle arrest and apoptosis

While most of the work to date support a pro-oncogenic and prometastatic role for GIV (Garcia-Marcos, Ghosh et al. 2010, Ghosh, Beas et al. 2010, Choi, Kim et al. 2017, Yang, Yang et al. 2018, Wang, Chen et al. 2020), a few have revealed a tumor suppressive function for GIV (Wang, Lei et al. 2018, Biehler, Wang et al. 2020). The pro-oncogenic roles have been demonstrated in stromal cells (Cos7, Vero) or epithelial cells with no (e.g., MDA MB231) or weak junctions (HeLa), whereas tumor suppressive effects were shown in cells with junctions (MDCK, Caco2, etc.)(Aznar, Patel et al. 2016, Biehler, Wang et al. 2020). Because the well-differentiated CRC cell line, Caco-2 are known to form well-defined junctions, was recently shown to have morphogenesis defects upon GIV-depletion (Biehler, Wang et al. 2020), and we confirmed that they express GIV-L **(Figure 2C)**, we used the control or the GIV-depleted (by shRNA) versions of this cell line as a model system for a series of phenotypic studies. First, we found that loss of GIV was associated also with a loss of contact-dependent growth arrest, as determined by growth of cells in patches of ‘piled-up’ cells in monolayers (circle; **Figure 7A)**. Second, the same cells produced colonies in larger number and size in anchorage-dependent colony growth assays **(Figure 7B-D)**. When the stained colonies were observed under light microscopy, colonies from GIV depleted cells were denser and more comprised of “piled up” cells (arrowheads; **Figure 7C)**, which is in keeping with our observations in monolayers **(Figure 7A)**. Third, higher colony growth was associated with a higher metabolic activity, as determined by the enhanced ability of GIV-depleted cells to metabolize the tetrazolium dye, MTT, irrespective of confluency **(Figure 7E)**.

**Figure 7.**
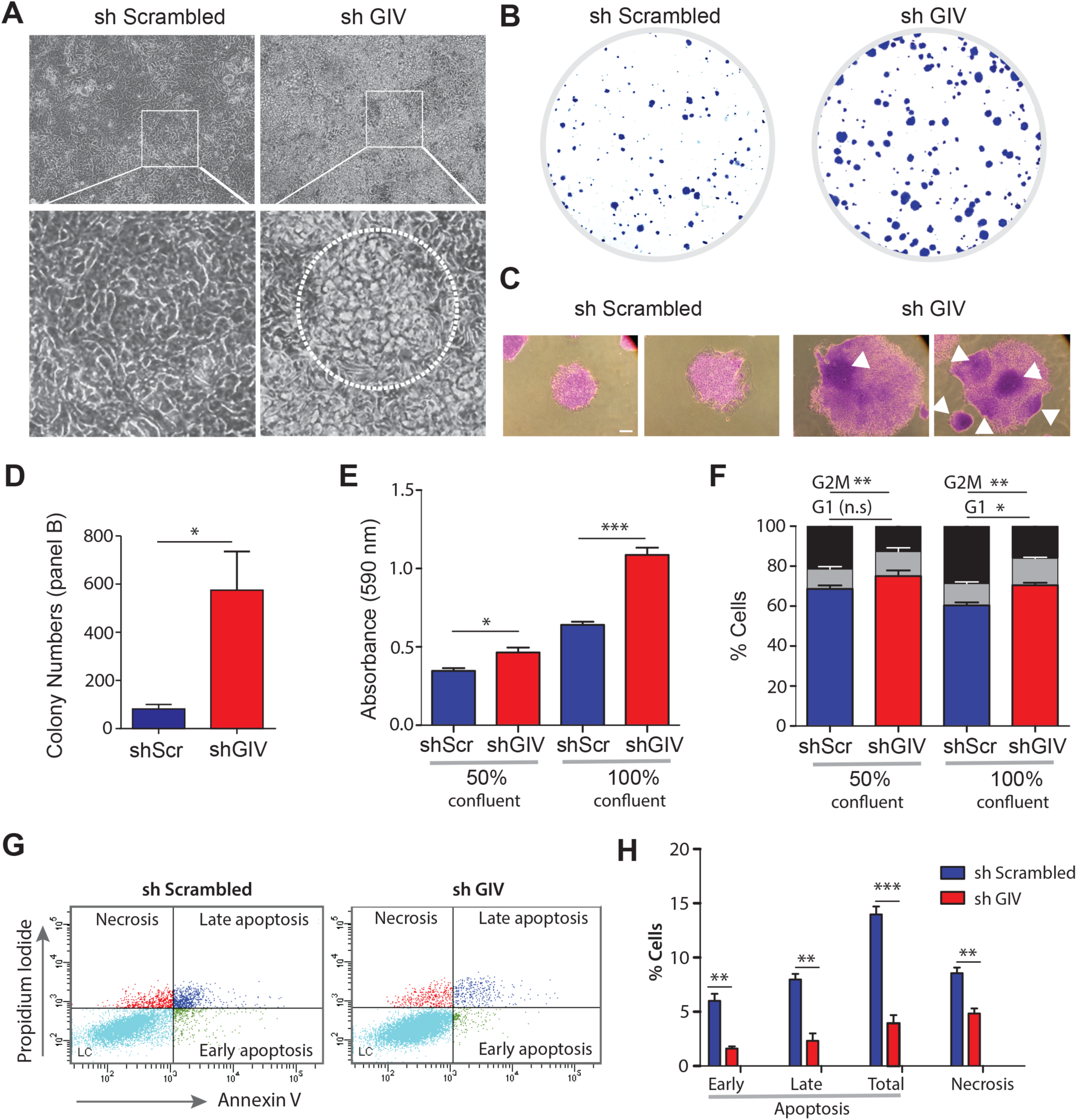
Depletion of GIV in Caco-2 cells increases anchorage-dependent colony growth, survival, loss of contact-dependent cell-cycle inhibition and reduced cell death. **A)** Phase contrast microscopy images of Caco-2 cells stably expressing a Sh Scrambled or Sh GIV construct. Caco-2 cells were cultured and grown in a confluent monolayer state for 10 days. Central ‘piling up’ of cells is frequently observed in the shGIV monolayer (as outlined). **B-D)** Representative images of crystal violet stained colonies, as seen during anchorage-dependent colony growth assays on control (sh Scrambled) and GIV-depleted (sh GIV) Caco-2 cells after 14 days in culture. Light microscopy images of representative colonies in C show the dense areas of piled up cells in sh GIV Caco-2 colonies (arrowheads). Bar graphs (D) show quantification of colonies. Error bars represent S.E.M; n = 3 (*) indicates p ≤0.05, as determined by student’s t-test. **E)** MTT proliferation assay on control (sh Scrambled) and GIV-depleted (sh GIV) Caco-2 cells grown at 50% or 100% confluency. Bar graphs show quantification of absorbance at 590 nm. Error bars represent S.E.M; n = 3. (*\**) indicates p<0.05, and (***) indicates p<0.001, as determined by student’s t-test. **F)** Cell cycle distribution of control (sh Scrambled) and GIV-depleted (sh GIV) Caco-2 cells grown at 50% or 100% confluency. Bar graphs show % of cells in each phase of the cell cycle. Error bars represent S.E.M; n = 3. (*) indicates p<0.05, (**) indicates p<0.01, n.s. non-significant, as determined by student’s t-test. **G-H)** Representative cytograms (G) of apoptotic and necrotic control (sh Scrambled) and GIV-depleted (sh GIV) Caco-2. The lower-right (annexin V^+^PI^-^ cells) and the upper-right (annexin V^+^PI^+^ cells) quadrants show early and late apoptotic cells, respectively, while the lower-left (annexin V^-^PI^-^ cells) and the upper-left (annexin V^-^PI^-^ cells) quadrants represent viable and necrotic cells, respectively. H) Bar graphs display the % of apoptotic and necrotic cells in panel G. Error bars represent S.E.M; n = 3. **P<0.01, ***P<0.001, as determined by student’s t-test.

To determine if the observed higher growth in GIV-depleted cells was due to merely higher proliferation, or lower cell death or both, we assessed for the distribution of cells across different stages of the cell cycle **(Figure 7F)** and for the population of cells undergoing cell death **(Figure 7G-H)**. We observed an increase in G0/G1 phase is observed (albeit only significantly observed at 100% confluency) in GIV depleted cells. This increase was accompanied with a concomitant decrease in the distribution of cells in the G2/M phase. When cell death was analyzed under the same conditions, we observed an overall decrease in cell death in GIV depleted cells (**Figure 7G-H**). Together, these findings highlight GIV’s tumor suppressive role in the well-differentiated Caco-2 cell line.

### GIV-L, but not GIV is suppressed during normal to adenoma progression in the colon

The increase in cell proliferative properties prompted us to examine the expression of GIV and GIV-L in normal and adenoma tissues **(Figure 8A-B)**. To this end, we utilized custom made antibodies raised against unique epitopes on GIV or GIV-L **(Figure S4A)**. Validation studies confirmed their specificity and ruled out cross reactivity against isoforms **(Figure S4B)**. Furthermore, they were tested alongside several commercially available antibodies that are expected to detect both isoform (total GIV). In normal healthy colonic tissue **(Figure 8A** and **C)**, we observed total GIV expression, as determined by GIV (CC-Ab) antibody, ubiquitously throughout the colon epithelial layer. Interestingly, staining with the specific GIV (CT-Ab) and GIV-L (CT-Ab) showed that GIV (CT-Ab) is also ubiquitously expressed (both in cytosolic and nuclear staining); however, GIV-L (CT-Ab) is restricted to the surface epithelial layer. In matched adenoma tissues **(Figure 8B-C)**, we observed an increase in GIV (CT-Ab) levels, but this increase was restricted to nuclei expression. No change in the cytosolic pool was observed. With regards to GIV-L (CT-Ab) staining, we observed a decrease in expression.

**Figure 8.**
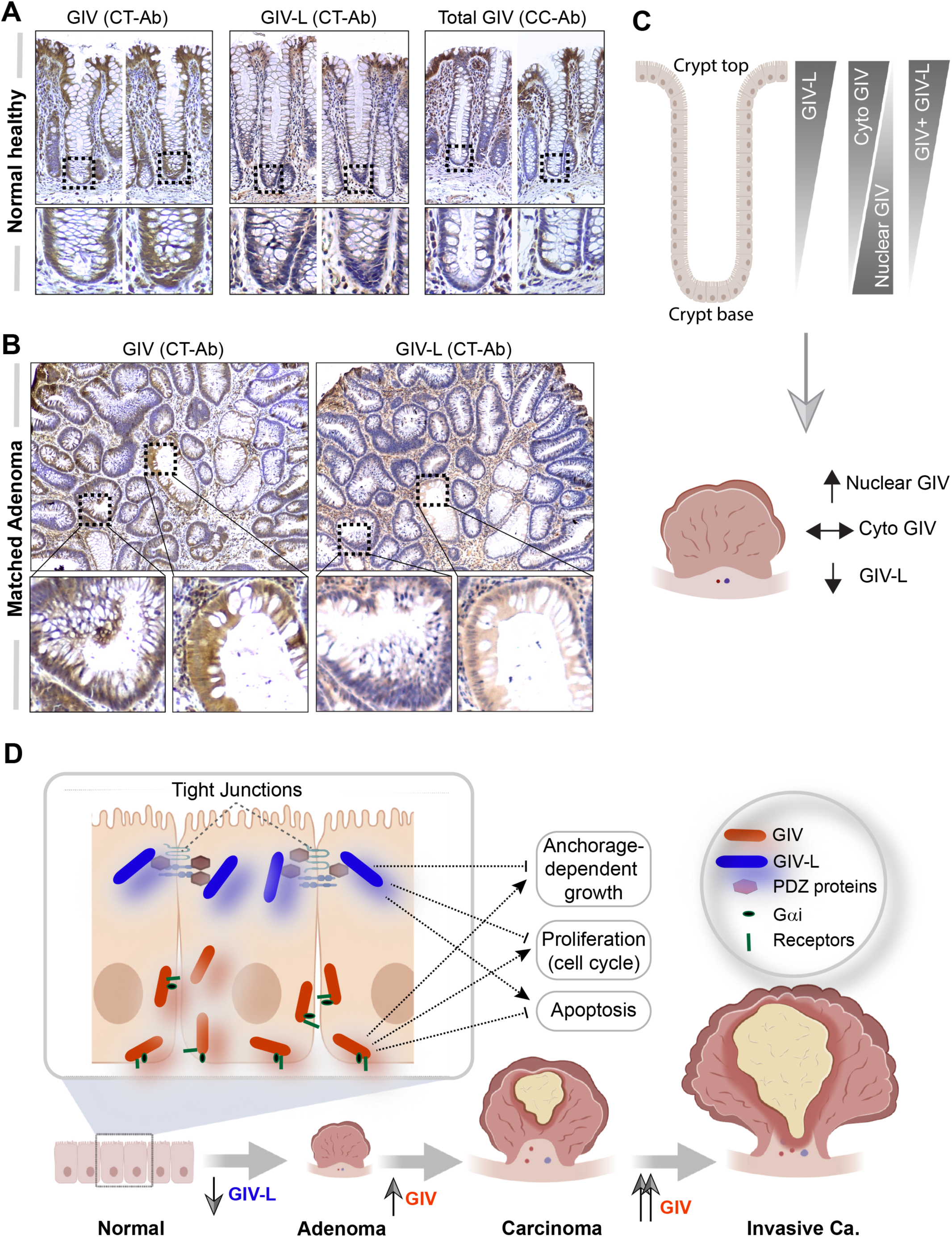
GIV-L is preferentially expressed in the surface epithelium of colon crypts, and is downregulated in the transformed epithelium in colon polyps. **A-B)** Images representative of patters of GIV staining, as determined by immunohistochemistry staining on normal healthy human colon (A) and matched adjacent adenoma (B) with various GIV (total and isoform specific) antibodies. See also *Supplemental Figure 3* for validation studies on the antibodies. **C)** Schematic summarizing the observed expression pattern observed in panels A (top) and B (bottom). ↑, ↓ and ↔ indicates upregulation, downregulation, and no discernible changes in expression, respectively. **D-E)** Working model of the opposing roles (D) and patterns of altered expression (E) of GIV and GIV-L isoforms in the colonic epithelium. Cytosolic GIV promotes stemness, growth, survival and cell migration whereas, cell-junction localized GIV-L inhibits growth, survival cell cycle, cell death.

These histological observations, taken together with those from our cell-based models, lead us to propose the following working model (see legend; **Figure 8D)** in which GIV and GIV-L perform opposing functions to maintain epithelial homeostasis, and that such functions are dictated based on protein subcellular localization. When GIV is cytosolic, it couples readily with receptors on the basolateral surface and with G proteins to primarily enhance signals that promote stemness, growth, survival and cell migration/invasion. By contrast, GIV-L is on cell-junction, where it senses junctional integrity and signals to inhibit growth, cell cycle progression and instead, promote cell death. What those signaling pathways are, remain to be determined.

## Discussion

### CCDC88A (GIV/Girdin) shows evolutionary flexibility of modularity

In this work, we confirmed the presence of two isoforms of GIV in vertebrates; besides the isoform that was known to exist and participate in the finetuning of endomembrane trimeric GTPase signaling downstream of multiple receptors, we show that there exists another long isoform of GIV, GIV-L, which contains a PBM. The absence or presence of the PBM determined GIV’s localization to cell-cell junctions, interactions with PDZ proteins, ability to bind and dissociate Gi trimers using its GEM motif and GIV’s overall functions in junction-containing epithelial cells. That the PBM is also present in invertebrates indicates that the motif has been conserved across evolution and not lost, as previously hypothesized (Nechipurenko, Olivier-Mason et al. 2016), suggesting that the PDZ-interacting module on GIV appeared early and remained conserved throughout evolution. By contrast, the GEM motif is absent in invertebrates, but present across all vertebrate species studied, suggesting that the G protein regulatory module on GIV appeared later in evolution. The zebrafish form of GIV (zGIV) contain both a functional GEM and PBM motif, representing the earliest species in which both modules coexist in the same protein. Furthermore, expression of zGIV is found on lateral line hair cells, a specialized ciliated cell (Metcalfe, Kimmel et al. 1985, McHenry and van Netten 2007). Given the well-established importance of G-proteins and Dvl in cilia function and positioning (Park, Mitchell et al. 2008, Ezan, Lasvaux et al. 2013, Mykytyn and Askwith 2017, Nachury and Mick 2019), it is plausible that zGIV regulates G-protein and PDZ interaction in these polarized epithelial cells.

### Modularity dictates localization, interactomes and function

We also confirmed GIV-L exists as a transcript and is translated into a functional protein also also in humans; in this case, it contains the PBM and approximately 150 extra amino acids. This longer isoform may have been previously missed for various reasons, including the absence of the transcript in the cDNA library used to identify GIV or due to an incomplete annotation of transcripts at the time the gene was discovered (Enomoto, Murakami et al. 2005, Le-Niculescu, Niesman et al. 2005). Prior work has shown that GIV localizes to various sub-cellular compartments, including on cell junctions of mammalian cells (Aznar, Midde et al. 2015). Furthermore, GIV was found to interact with PDZ proteins such as Dvl and ParD3 (Sasaki, Kakuwa et al. 2015, Boldt, van Reeuwijk et al. 2016), although how such interactions could be mediated remained unsolved. Cell junctions are known to be clustered with PDZ proteins where a complex PDZ-PBM interaction network regulates signaling (Fanning and Anderson 1999, Subbaiah, Kranjec et al. 2011, Manjunath, Ramanujam et al. 2018). By showing that GIV-L, and not GIV or a GIV-L mutant that lacks the PBM, localizes to cell junctions and binds to PDZ proteins, we show that both localization and PDZ-binding requires GIV-L’s PBM. Furthermore, we found that GIV-L transcript was readily detectable in some cell lines, but not others; expression was virtually restricted to epithelial lines that readily make cell-cell junctions. We conclude that between the two isoforms of GIV that co-exist in cells, it is GIV-L that finetunes junctional signaling through its PBM.

Among the phenotypic readouts investigated here, we focused on key epithelial phenotypes previously studied in the context of GIV. Others and us have documented on numerous instances that mammalian GIV supports cellular phenotypes, e.g., cell proliferation, growth, survival, migration and invasion (Enomoto, Murakami et al. 2005, Garcia-Marcos, Ghosh et al. 2010, Ohara, Enomoto et al. 2012, Choi, Kim et al. 2017, Yang, Yang et al. 2018, Wang, Chen et al. 2020). In addition, numerous groups have documented that GIV expression goes up during neoplastic progression in numerous cancers (reviewed in (Ghosh 2015)). Based on these observations, GIV has been generally believed to serve primarily as an oncogene that fuels cancer initiation and progression. However, this belief has recently been challenged by others who have suggested a tumor suppressive role for GIV (Biehler, Wang et al. 2020), although the molecular mechanism(s) for such contrasting roles for the same protein was not revealed. By showing here that GIV-L maintains epithelial homeostatic properties (e.g., junction-dependent cell cycle and growth inhibition and apoptosis), we provide the first insights into how GIV may perform opposing roles in epithelial cells via its two isoforms. Because cells junctions, in addition to its adhesive properties, are well-regarded as a cellular structure that blocks tumor growth (Subbaiah, Kranjec et al. 2011), GIV maintains epithelial integrity (Sasaki, Kakuwa et al. 2015, Aznar, Patel et al. 2016, Biehler, Wang et al. 2020), we propose a working model (see **Figure 8D**) where the two opposing functions of GIV, i.e., tumor suppressor vs. oncogene are driven by its subcellular localizations; localization at cell junctions enables GIV-L to exert its tumor suppressive functions whereas localization in cytosol enables GIV to access various receptors and G proteins on basolateral membranes to exert its pro-oncogenic signaling functions. This model is further supported by the detection of GIV-L transcript in colorectal cancer cells that form cell junctions and a lack of transcripts in cells that do not form junctions. That the expression of GIV-L, but not GIV is suppressed in colon tissue during adenoma formation further supports the model that GIV-L is likely to be the tumor-suppressive isoform.

### CCDC88 proteins exemplify evolutionary enrichment of how PBMs may finetune G protein/receptor signaling

It is noteworthy that CCDC88A/GIV is not the only member of the CCDC88 family that has a PBM. CCDC88C/Daple also features a C-terminal PBM motif that is similar but not identical in sequence to that in GIV-L, raising the possibility that GIV-L has a unique PDZ interactome. Analysis of our own BioID proximity labeling studies, and others, reveal that GIV-L indeed binds to several PDZ domain-containing proteins, which only partially overlaps with that of Daple (Ear, Saklecha et al. 2020). Interaction assays confirmed that GIV-L and Daple can indeed bind to the some common PDZ-proteins (e.g., ParD3 and Dvl).

We also provided evidence for how the PBM•PDZ interactions of members of the CCDC88 family may represent a mechanism *via* which G protein signaling *via* GEM motifs in CCDC88 appear to have been subjected to higher orders of regulatory controls during evolution. For example, by showing that GIV-L bound G protein Gαi preferably in the presence of Dvl, we demonstrated modular cooperativity. These findings add to the prior examples of competition and co-existence between PBM•PDZ and GEM•G protein interactions in CCDC88C (Aznar, Ear et al. 2018, Ear, Saklecha et al. 2020). Why some PDZs cooperate and/or co-exist in complexes with CCDC88 and Gαi proteins, while others compete remains unknown and warrants further investigation.

In conclusion, we have identified a novel isoform of GIV that contains an evolutionarily conserved PBM and demonstrate how evolutionary flexibility between two isoforms of GIV—one without and one with PBM-dictates protein localization, interactome and functions. Insights into how binding to PDZ proteins shape GIV’s localization and interactions have revealed how modularity regulates GIV’s functions. Such revelation can help us to further understand the role GIV plays in tissue homeostasis and how its dysregulation may trigger diseases.

## ACKNOWLEDGEMENTS

This paper was supported by NIH CA238042, AI141630, CA100768 and CA160911 as well as UG3TR002968 (to P.G). J.E was supported by an NCI/NIH-funded Cancer Biology, Informatics & Omics (CBIO) Training Program (T32 CA067754) and a Postdoctoral Fellowship from the American Cancer Society (PF-18-101-01-CSM). S.K and D.S are supported by R01GM138385 (to D.S).

## AUTHOR CONTRIBUTIONS

J.E, A. A. A.E-H, N.R, J.C. and P.G designed, executed and analyzed most of the experiments in this work. S.R. and I.K designed, executed and analyzed the BRET assays. T.N. carried out the analyses of the HCM with supervision from I.K. S.K and D.S. executed and analyzed the IHC studies. J.E, A. A. A.E-H, and S.R wrote methods. J.E and P.G conceived the project, wrote and edited the manuscript.

## Materials and Methods

- KEY RESOURCE TABLE
- CONTACT FOR REAGENT AND RESOURCE SHARING
- EXPERIMENTAL MODEL AND SUBJECT DETAILS

- Human IHC Samples
- Cell Lines (DLD1, HCT116, Caco-2, HEK293T, HeLa)
- Zebrafish (WT-AB)
- METHOD DETAILS

- Immunofluorescence and Confocal Microscopy, Image analysis
- Image Processing
- Cell Fractionation
- Immunoblotting
- GIV CRISPR/Cas9 Gene Editing and Validation
- Anchorage-dependent and Anchorage-independent Colony Growth Assays
- Biotin Proximity Labeling
- In Gel Digest
- LC-MS analysis
- Gene Ontology analysis
- Recombinant Protein Purification
- In Vitro GST-Pulldown and In-cellulo Co-immunoprecipitation (CoIP) Assays
- RNA probe synthesis & Whole-mount RNA in situ hybridization
- Gi-directed response assay & Bioluminescence resonance energy transfer (BRET) experiment
- Cell cycle and apoptosis assay
- Immunohistochemistry
- QUANTIFICATION AND STATISTICAL ANALYSIS

- Statistical Analysis

## Key Resource Table

| REAGENT or RESOURCE | SOURCE | IDENTIFIER |
| --- | --- | --- |
| **Antibodies** |  |  |
| Rabbit polyclonal anti-Gαi3 (C-10) | Santa Cruz Biotechnology | N/A |
| Rabbit polyclonal anti-ZO1 | GeneTex | GTX108613 |
| Rabbit polyclonal anti-ZO1 | GeneTex | GTX108627 |
| Rabbit polyclonal anti-PARD3 | Proteintech | 11085-1-AP |
| Rabbit polyclonal anti-E-cadherin | Santa Cruz Biotechnology | sc-7870 |
| Rabbit polyclonal anti-β-tubulin | Santa Cruz Biotechnology | sc-9104 |
| Rabbit polyclonal anti-GIV (Girdin) (T-13) | Santa Cruz Biotechnology | sc-133371 |
| Rabbit polyclonal anti-GIV (Girdin) (CC-Ab) | Millipore Sigma | ABT80 |
| Rabbit polyclonal anti-GIV (CT-Ab) | *This paper* | N/A |
| Rabbit polyclonal anti-GIV-L (CT-Ab) | *This paper* | N/A |
| Mouse monoclonal anti-GIV (Girdin) (H-6) | Santa Cruz Biotechnology | sc-393757 |
| Mouse monoclonal anti-β-catenin | Santa Cruz Biotechnology | sc-7963 |
| Mouse monoclonal anti-GAPDH | Santa Cruz Biotechnology | sc-365062 |
| Mouse monoclonal anti-αE-catenin | Santa Cruz Biotechnology | sc-9988 |
| Mouse monoclonal anti-FLAG | Millipore Sigma | MAB3118 |
| Mouse monoclonal anti-FLAG (hybridoma) | Purified in house | N/A |
| Mouse monoclonal anti-GST | GenScript | A00865 |
| Mouse monoclonal anti-Myc | Cell Signaling Technology | 2276S |
| Mouse monoclonal anti-Myc (hybridoma) | Purified in house | N/A |
| Mouse monoclonal anti-Dvl | Santa Cruz Biotechnology | sc-166303 |
| Mouse monoclonal anti-α-tubulin | Santa Cruz Biotechnology | sc-5286 |
| Mouse monoclonal anti-EGFP | Santa Cruz Biotechnology | Sc-9996 |
| Goat anti-Rabbit IgG (680) | LI-COR Biosciences | 926-68071 |
| Goat anti-Rabbit IgG, Alexa Fluor 594 conjugated | ThermoFisher Scientific | A11072 |
| Goat anti-Mouse IgG (800) | LI-COR Biosciences | 926-32210 |
| Goat anti-Mouse IgG, Alexa Fluor 488 conjugated | ThermoFisher Scientific | A11017 |
| ImmPRESS® HRP Horse Anti-Rabbit IgG Polymer Detection Kit, Peroxidase | Vector Laboratories | MP-7401 |
| Anti-Digoxigenin-AP | Roche | 11093274910 |
| **Biological Samples and Cell Lines** |  |  |
| DLD1 | ATCC | CCL-221 |
| HCT116 | ATCC | CCL-247 |
| Caco-2 | ATCC | HTB-37 |
| HeLa | ATCC | CCL-2 |
| HEK293T | ATCC | CRL-11268 |
| Caco-2 ShGIV | Biehler, et. al., 2020 | N/A |
| Caco-2 ShScrambled | Biehler, et. al., 2020 | N/A |
| **Chemicals, Kits, Recombinant Proteins, and Plasmids** |  |  |
| Streptavidin, Alexa Fluor® 680 conjugate | ThermoFisher Scientific | S21378 |
| Streptavidin, Alexa Fluor® 594 conjugate | ThermoFisher Scientific | S11227 |
| Streptavidin Magnetic Beads | ThermoFisher Scientific | 88816 |
| Biotin | Sigma-Aldrich | B4639-500MG |
| Coelenterazine-h | Promega | S2011 |
| DIG RNA Labeling Mix | Roche | 11277073910 |
| T7 RNA polymerase | Promega | P2075 |
| MTT | Millipore Sigma | 475989-1GM |
| Guava Cell Cycle Reagent | Millipore Sigma | 4700-0160 |
| ImmPACT® DAB Substrate, Peroxidase (HRP) | Vector Laboratories | SK-4105 |
| Dead Cell Apoptosis Kit with Annexin V Alexa Fluor™ 488 & Propidium Iodide (PI) | ThermoFisher Scientific | V13241 |
| DAPI (4',6-Diamidino-2-Phenylindole, Dilactate) | Thermo Fisher Scientific | D3571 |
| pSpCas9(BB)-2A-Puro (PX459) V2.0 | Addgene | 62988 |
| pcsDest2-GST | *This paper* | N/A |
| pcsDest2-GST GIV-L CT (1660-2026) | *This paper* | N/A |
| pcsDest2-EGFP | *This paper* | N/A |
| pcsDest2-EGFP-zGIV-CT | *This paper* | N/A |
| pcsDest2-EGFP-zDaple-CT | *This paper* | N/A |
| myc-pcDNA 3.1 (+) – GIV (full length) | *This paper* | N/A |
| myc-pcDNA 3.1 (+) – GIV-F1685A (full length) | *This paper* | N/A |
| myc-pcDNA 3.1 (+) – GIV-L (full length) | *This paper* | N/A |
| myc-pcDNA 3.1 (+) – GIV-L-F1685A (full length) | *This paper* | N/A |
| myc-pcDNA 3.1 (+) – GIV-L-ΔPBM (full length) | *This paper* | N/A |
| myc-pcDNA 3.1 (+) – GIV-L-F1685A/ΔPBM (full length) | *This paper* | N/A |
| pcDNA3.1-N-term FLAG-PARD3 (full length) | Peng Zhang, et. al., 2016 | N/A |
| pcDNA3.1-N-term FLAG-PARD3 (ΔPDZ 1) | Peng Zhang, et. al., 2016 | N/A |
| pcDNA3.1-N-term FLAG-PARD3 (ΔPDZ 2) | Peng Zhang, et. al., 2016 | N/A |
| pcDNA3.1-N-term FLAG-PARD3 (ΔPDZ 3) | Peng Zhang, et. al., 2016 | N/A |
| pcDNA3.1(+)-hGai1(91)-RLuc2 | Brown et al. 2010 | N/A |
| mVenus-hGBB1 | Brown et al. 2016 | N/A |
| mVenus-hGBG2 | Brown et al. 2016 | N/A |
| **Software** |  |  |
| ImageJ | National Institute of Health | https://imagej.net/Welcome |
| DAVID 6.8 | DAVID Bioinformatics Resources | https://david.ncifcrf.gov/home.jsp |
| Molsoft ICM v3.8-6 | Molsoft LLC | http://www.molsoft.com/index.html |

### Cell lines and culture methods

DLD1 cells were cultured using RMPI media containing 10% FBS. Cells were routinely passaged at a dilution of 1:5 to 1:10. HEK293T, Caco-2, and HeLa cells were cultured using DMEM media containing 10% FBS and routinely passaged at a dilution of 1:5 to 1:10. HCT116 cells were cultured in McCoy’s 5a Medium containing 10% FBA and passed at a dilution of 1:5 to 1:10. Caco-2 (Sh Scrambled and Sh GIV) cultured as previously described (Biehler, Wang et al. 2020).

### Zebrafish Husbandry

Zebrafish protocol and maintenance was performed using methods approved by the University of California, San Diego Institutional Animal Care and Use Committee (UCSD-IACUC). Zebrafish wild type (AB) strains were used for tissue expression studies.

### RNA probe synthesis & whole-mount RNA in situ hybridization

DNA template used in the in vitro transcription of RNA probes was amplified from a pooled cDNA library of zebrafish embryos (12 to 72 hours post fertilization). Flanking the reverse primer was a T7 RNA polymerase site. PCR amplicon was separated on an agarose gel and then extracted using a DNA gel extraction kit (Zymo). Purified DNA was used in an in vitro transcription reaction using T7 RNA polymerase (Promega) and DIG RNA labeling mix (Roche). Transcription was carried out at 37°C for two hours, followed by the addition of DNase to degrade the template. EDTA was added to ensure a full stop of the transcription reaction. mRNA probe was purified using lithium chloride precipitation and diluted in hybridization buffer (50% formamide, 750mM NaCl, 75mM Sodium Citrate, 50μg/ml Heparin, 5 mM EDTA, 0.5mg/ml rRNA, 1 mM Citric Acid, 0.1% Tween-20, pH=6.0) prior to use in whole-mount RNA in situ hybridization.

Whole mount *in situ* hybridization was performed using the labeled probed as previously described (Jowett 1999). Briefly, embryos were first fixed overnight at 4°C using 4%-paraformaldehyde (PFA) in PBS. After fixation, PFA was washed four times (15 min each) with PBS, followed by two washes (5 min each) with 100% Methanol. At this point, bleaching was performed to remove pigmentation, if necessary. Embryos were rehydrated using successive washes of PBT (0.2% BSA and 0.2% Tween-20 in 1X PBS). Embryos were then treated with proteinase K, washed using PBT, and then equilibrated into hybridization buffer at 65°C for 2 hours. Afterwards, embryos were when transferred to hybridization buffer containing DIG-labeled RNA probe and allowed to incubated overnight at 65°C. Excess probed was removed through excessive washes using hybridization buffer, followed by a gradient of SSC solution (150 mM NaCl, 15 mM Sodium Citrate) and PBT buffer washes. Incubation with anti-DIG secondary antibody was carried out overnight at 4°C followed by development using Nitro Blue Tetrazolium (NBT) and 5-bromo-4-chloro-3-indolyl-phosphate (BCIP). Alkaline-phosphatase reaction was stopped through excessive washes using PBT and embryos were immediately mounted on methylcellulose and imaged using a light-microscope.

### Recombinant Protein Purification

GST-tagged proteins were expressed in E. coli stain BL21 (DE3) and purified. Cultures were induced using 1mM IPTG overnight at 25°C. Cells were then pelleted and resuspended in GST lysis buffer (25 mM Tris-HCl, pH 7.5, 20 mM NaCl, 1 mM EDTA, 20% (vol/vol) glycerol, 1% (vol/vol) Triton X-100, 2×protease inhibitor cocktail). Cells were lysed by sonication, and lysates were cleared by centrifugation at 12,000 X g at 4°C for 30 mins. Supernatant was then affinity purified using glutathione-Sepharose 4B beads (GE Healthcare), followed by elution, overnight dialysis in PBS, aliquoted and then stored at -80°C.

### In Vitro GST-Pulldown and In-cellulo Co-immunoprecipitation (CoIP) Assays

Purified GST-tagged proteins from E. coli were immobilized onto glutathione-Sepharose beads and incubated with binding buffer (50 mM Tris-HCl (pH 7.4), 100 mM NaCl, 0.4% (v:v) Nonidet P-40, 10 mM MgCl_2_, 5 mM EDTA, 2 mM DTT) for 60 mins at room temperature. For the pulldown of protein-protein complexes from cell lysates, cells were first lysed in cell lysis buffer (20 mM HEPES, pH 7.2, 5 mM Mg-acetate, 125 mM K-acetate, 0.4% Triton X-100, 1 mM DTT, 500 μM sodium orthovanadate, phosphatase inhibitor cocktail (Sigma-Aldrich) and protease inhibitor cocktail (Roche)) using a 28G needle and syringe, followed by centrifugation at 10,000Xg for 10 mins. Cleared supernatant was then used in binding reaction with immobilized GST-proteins for 4 hours at 4°C. After binding, bound complexes were washed four times with 1 ml phosphate wash buffer (4.3 mM Na2HPO4, 1.4 mM KH2PO4, pH 7.4, 137 mM NaCl, 2.7 mM KCl, 0.1% (v:v) Tween 20, 10 mM MgCl_2_, 5 mM EDTA, 2 mM DTT, 0.5 mM sodium orthovanadate). Bound proteins were then eluted through boiling at 100°C in sample buffer.

For CoIP assays, cells lysates (as prepared above) was incubated with capture antibodies for 3 hours at 4°C, followed by the addition of Protein A or Protein G beads to capture antibody bound protein-protein complexes. Bound proteins were then eluted through boiling at 100°C in sample buffer.

### BRET-based assessment of Gai/Gβγ dissociation

mVenusCT-hGBB1 and mVenusNT-hGBG2 were a gift from Nevin Lambert, Univ. of Alberta (Brown, Lambert et al. 2016). pcDNA3.1(+)-hGai1(91)-RLuc2 was generously shared by Michel Bouvier (Breton, Sauvageau et al. 2010).

On day 1, HEK293T cells were plated at a density of 3.5×10^5^ cells per well into a 12-well plate using DMEM containing 10%FBS. On day 2, cell culture media was replaced with fresh media and then cells were transfected with 0.2 μg/well CXCR4, 0.2 μg/well VenusCT-Gβ, 0.2 μg/well VenusNT-Gγ, 20 ng/well Gai(91)-Rluc2, and 0.4 μg/well of GIV variants or pcDNA vector control using TransIT-X2 transfection reagent (MirusBio) according to manufacturer’s instruction. On day 3, cells were lifted by pipetting and transferred into 1.5 mL microcentrifuge tubes, spun down, and resuspended in DMEM+10% FBS to 40,000 cells/mL. 100 uL of the cell suspension was then replated on a poly-D-lysine-coated 96-well black/clear bottom plate and allowed to adhere. On day 4, cell culture media was carefully removed and replaced with 80 uL of serum-free assay buffer (PBS + 0.1% glucose) for 60 minutes. The luciferase substrate, coelenterazine-h (10 uM final), was added to each well. The plate was incubated at room temperature for 5 min, after which repeated readings of light emission at 485 and 515 nm were initiated using the Victor X luminescence plate reader (Perkin Elmer, USA) over the course of 3 min. The average BRET was calculated over 3 min after adding Coelenterazine-h. The experiment was repeated in three independent biological replicates on different days, each containing three technical replicates. An average of the three biological replicate is shown, and graphs were plotted using GraphPad Prism 5.

### Cell Fractionation

Cells were harvested and suspended in homogenization buffer (10 mM sodium phosphate buffer [pH 7.2], 1 mM MgCl_2_, 30 mM NaCl, 1 mM DTT, and 0.5 mM phenylmethylsulfonyl fluoride [PMSF], supplemented with protease and phosphatase inhibitors), and homogenized using a 30-gauge needle and syringe. Unlysed cells were cleared by centrifugation at 1,000 × *g* for 10 min at 4°C and collecting supernatant. Crude membranes were separated from the homogenate by centrifugation of post-nuclear supernatant at 100,000 × *g* for 60 min at 4°C in a TLA-41 fixed-angle rotor in a TLA-100 table-top ultracentrifuge (Beckman Coulter, Krefeld, Germany). Pelleted membranes were washed in homogenization buffer before resuspension in cell lysis buffer containing 0.4% Tx-100.

### Quantitative immunoblotting

For immunoblotting, protein samples were boiled in Laemmli sample buffer, separated by SDS-PAGE and transferred onto 0.4μm PVDF membrane (Millipore) prior to blotting. Post transfer, membranes were blocked using 5% Non-fat milk or 5% BSA dissolved in PBS. Primary antibodies were prepared in blocking buffer containing 0.1% Tween-20 and incubated with blots, rocking overnight at 4°C. After incubation, blots were incubated with secondary antibodies for one hour at room temperature, washed, and imaged using a dual-color Li-Cor Odyssey imaging system.

### Immunofluorescence and Confocal Microscopy, Image analysis

Cells were fixed using -20°C methanol (or 4°C paraformaldehyde, PFA) for 20 to 30 mins, rinse with PBS, then permeabilized for 1hr using blocking/permeabilization buffer (0.4% Triton X-100 and 2 mg/ml BSA dissolved in PBS). Primary antibody and secondary antibody were diluted in blocking buffer and incubated with cells for 1 hr each. Coverslips were mounted using Prolong Gold (Invitrogen) and imaged using a Leica SPE CTR4000 confocal microscope.

### Image Processing

All images were processed on ImageJ software (NIH) and assembled into figure panels using Photoshop and Illustrator (Adobe Creative Cloud). Some images were created using BioRender.com. All graphs were generated using Excel (Microsoft) or GraphPad Prism.

### GIV CRISPR/Cas9 Gene Editing and Validation

GIV guide DNA sequence was cloned into PX-459 vector and transfected into cells using PEI. For selection, puromycin was added to cells and when untransfected control plates showed 95 to 100% cell death, cells were washed with PBS and media (without puromycin) was added to cells for 8 hours. Following recovery, cells were resuspended and sparsely plated (approximately 30 cells/plate) onto 10 cm plates so that individual cell colonies could be isolated and picked into 12-well plates for screening.

To identify cell clones harboring mutations in gene coding sequence, genomic DNA was extracted using 50 mM NaOH and boiling at 95°C for 60mins. After extraction, pH was neutralized by the addition of 10% volume 1.0 M Tris-pH 8.0. The crude genomic extract was then used in PCR reactions with primers flanking the targeted site. Amplicons were analyzed for insertions/deletions (indels) using a TBE-PAGE gel. Indel sequence was determined by cloning amplicons into a TOPO-TA cloning vector (Invitrogen) following manufacturer’s protocol.

### Biotin Proximity Labeling

BioID was performed as previously described (Ear, Saklecha et al. 2020). Briefly, HEK293T were plated 24 hrs prior to transfection with mycBirA-tagged GIV construct. Thirty hours post transfection, cells were treated with 50 μM biotin (dissolved in culture media) for 16 hrs. Cells were then rinsed two times with PBS and lysed by resuspending in lysis buffer (50 mM Tris, pH 7.4, 500 mM NaCl, 0.4% SDS, 1 mM dithiothreitol, 2% Triton X-100, and 1× Complete protease inhibitor) and sonication in a bath sonicator. Cell lysates were then cleared by centrifugation at 20,000 X g for 20 mins and supernatant was then collected and incubated with streptavidin magnetic beads overnight at 4°C. After incubation, beads were washed twice with 2% SDS, once with wash buffer 1 (0.1% deoxycholate, 1% Triton X-100, 500 mM NaCl, 1 mM EDTA, and 50 mM HEPES, pH 7.5), followed with once wash using wash buffer 2 (250 mM LiCl, 0.5% NP-40, 0.5% deoxycholate, 1 mM EDTA, and 10 mM Tris, pH 8.0), and once with 50 mM Tris pH 8.0. Biotinylated complexes were then eluted using sample buffer containing excess biotin and heating at 100°C. Prior to mass spectrometry identification, eluted samples were run on SDS-PAGE and proteins were extracted by in gel digest.

### In Gel Digest

Protein digest and mass spectrometry was perform as previously described (Shevchenko, Wilm et al. 1996). Briefly, the gel slices were cut into 1mm × 1 mm cubes, destained 3 times by first washing with 100 ul of 100 mM ammonium bicarbonate for 15 minutes, followed by the addition of equal volume acetonitrile (ACN) for 15 minutes. The supernatant was collected, and samples were dried using a speedvac. Samples were then reduced by mixing with 200 µl of 100 mM ammonium bicarbonate-10 mM DTT and incubated at 56°C for 30 minutes. The liquid was removed and 200 ul of 100 mM ammonium bicarbonate-55mM iodoacetamide was added to gel pieces and incubated covered at room temperature for 20 minutes. After the removal of the supernatant and one wash with 100 mM ammonium bicarbonate for 15 minutes, equal volume of ACN was added to dehydrate the gel pieces. The solution was then removed, and samples were dried in a SpeedVac. For digestion, enough solution of ice-cold trypsin (0.01 ug/ul) in 50 mM ammonium bicarbonate was added to cover the gel pieces and set on ice for 30 min. After complete rehydration, the excess trypsin solution was removed, replaced with fresh 50 mM ammonium bicarbonate, and left overnight at 37°C. The peptides were extracted twice by the addition of 50 µl of 0.2% formic acid and 5 % ACN and vortex mixing at room temperature for 30 min. The supernatant was removed and saved. A total of 50 µl of 50% ACN-0.2% formic acid was added to the sample, and vortexed again at room temperature for 30 min. The supernatant was removed and combined with the supernatant from the first extraction. The combined extractions are analyzed directly by liquid chromatography (LC) in combination with tandem mass spectroscopy (MS/MS) using electrospray ionization.

### LC-MS analysis

Trypsin-digested peptides were analyzed by ultra-high-pressure liquid chromatography (UPLC) coupled with tandem mass spectroscopy (LC-MS/MS) using nano-spray ionization. The nanospray ionization experiments were performed using a Orbitrap fusion Lumos hybrid mass spectrometer (Thermo) interfaced with nano-scale reversed-phase UPLC (Thermo Dionex UltiMate™ 3000 RSLC nano System) using a 25 cm, 75-micron ID glass capillary packed with 1.7-µm C18 (130) BEH™ beads (Waters corporation). Peptides were eluted from the C18 column into the mass spectrometer using a linear gradient (5–80%) of ACN (Acetonitrile) at a flow rate of 375 μl/min for 1h. The buffers used to create the ACN gradient were: Buffer A (98% H_2_O, 2% ACN, 0.1% formic acid) and Buffer B (100% ACN, 0.1% formic acid). Mass spectrometer parameters are as follows; an MS1 survey scan using the orbitrap detector (mass range (m/z): 400-1500 (using quadrupole isolation), 120000 resolution setting, spray voltage of 2200 V, Ion transfer tube temperature of 275 C, AGC target of 400000, and maximum injection time of 50 ms) was followed by data dependent scans (top speed for most intense ions, with charge state set to only include +2-5 ions, and 5 second exclusion time, while selecting ions with minimal intensities of 50000 at in which the collision event was carried out in the high energy collision cell (HCD Collision Energy of 30%), and the fragment masses where analyzed in the ion trap mass analyzer (With ion trap scan rate of turbo, first mass m/z was 100, AGC Target 5000 and maximum injection time of 35ms). Protein identification and label free quantification was carried out using Peaks Studio 8.5 (Bioinformatics solutions Inc.)

### Gene Ontology Analysis

Proteins identified by mass spectrometry in biotin-treated samples, but not in non-biotin-treated samples were analyzed using DAVID. Functional annotation was grouped by INTERPRO protein domains for GO analysis. Classification with p-value less than 0.5 were considered as significant.

### Using Human Cell Map for the identification of potential interactors and subcellular compartment annotation of GIV

The HCM dataset was downloaded (accessed 01/06/2020), reprocessed, and rescored using SAINTexpress (Teo, Liu et al. 2014) with a modified negative control set only containing untransfected cells. Controls cells expressing cytoplasmic BirA*-FLAG or BirA*-FLAG-GFP (included in the original analysis (Go, Knight et al. 2019)) were removed. This was done to eliminate non-specific proteins that bound to streptavidin beads even in the absence of biotinylation, while retaining possible specific interactions in the cytoplasm. Following rescoring, a 1% Bayesian FDR cutoff was used to filter for confident bait-prey pairs and the resulting dataset was used to calculate prey-prey correlations with GIV, as per Go et al (Go, Knight et al. 2019). Annotation of protein subcellular localization was taken from the Non-negative Matrix Factorization method used in HCM (Go, Knight et al. 2019).

### Anchorage-dependent colony growth assay

Anchorage-dependent growth were monitored on regular tissue culture plastics by seeding cells at a density of 5,000 cells per well in a 6-well plate and incubation for approximately 21 days in 10% FBS media. Media was changed approximately every 3 days to ensure health of the cells. Cells were then fix and permeabilized using 100% methanol prior to staining with 0.1% crystal violet. Colony growth were imaged by light microscopy and colonies were counted using ImageJ (NIH).

### Cell cycle, apoptosis, and cell proliferation assay

Cell cycle analysis and apoptotic cell quantification was performed using the Guava cell cycle reagent (Millipore Sigma) or the annexin V/propidium iodide (PI) staining kit (Thermo Fisher Scientific), respectively, according to the manufacturer’s instructions. Cells were quantified on a BD LSR II flow cytometer and analyzed using FlowJo software (FlowJo, Ashland, OR, USA).

Cell proliferation was measured using the MTT reagent and cells cultured in 96-well plates. Cells were incubated with MTT for 4 hr at 37°C. After incubated, culture media was removed and 150μl of DMSO was added in order to solubilize the MTT formazan crystals. Optical density was determined at 590 nm using a TECAN plate reader. At least three independent experiments were performed.

### Immunohistochemistry

Slides containing normal colon and adjacent cancer tissue were deparaffinized in xylene and then rehydrated in a gradation of alcohols to water. Slides were immersed in sodium citrate buffer (pH 6.0) and pressure cooked for 3 minutes and 30 seconds for antigen retrieval. Endogenous peroxidase activity was blocked by incubation using hydrogen peroxide. To block non-specific protein binding, 2.5% goat serum was used. Tissues were then incubated with primary antibodies for one hour at room temperature and in a humidified chamber. Afterwards, slides were rinsed three times with PBS (5 minutes each rinse). Sections were then incubated with horse anti-rabbit HRP conjugated secondary antibody for 30 minutes at room temperature and then washed three times with PBS (5 minutes each rinse). Afterwards, development with DAB substrate and counterstain with hematoxylin. After development, slides were dehydrated in using a gradient of alcohol washes, cleared in xylene, and then mounted with coverslips. Epithelial and stromal components of tumors were identified by staining duplicate slides in parallel with hematoxylin and eosin and visualizing by light microscopy.

### Statistical Analysis and Replicates

Student’s t-test was used to determine significance with P values of < 0.05 set as the minimal threshold for statistical significance. Where statistical analysis was performed, experiments were performed (at least) in triplicates.

## References

Amacher, J. F., L. Brooks, T. H. Hampton and D. R. Madden (2020). “Specificity in PDZ-peptide interaction networks: Computational analysis and review.” J Struct Biol X 4: 100022.

Aznar, N., J. Ear, Y. Dunkel, N. Sun, K. Satterfield, F. He, N. A. Kalogriopoulos, I. Lopez-Sanchez, M. Ghassemian, D. Sahoo, I. Kufareva and P. Ghosh (2018). “Convergence of Wnt, growth factor, and heterotrimeric G protein signals on the guanine nucleotide exchange factor Daple.” Sci Signal 11(519).

Aznar, N., K. K. Midde, Y. Dunkel, I. Lopez-Sanchez, Y. Pavlova, A. Marivin, J. Barbazan, F. Murray, U. Nitsche, K. P. Janssen, K. Willert, A. Goel, M. Abal, M. Garcia-Marcos and P. Ghosh (2015). “Daple is a novel non-receptor GEF required for trimeric G protein activation in Wnt signaling.” Elife 4: e07091.

Aznar, N., A. Patel, C. C. Rohena, Y. Dunkel, L. P. Joosen, V. Taupin, I. Kufareva, M. G. Farquhar and P. Ghosh (2016). “AMP-activated protein kinase fortifies epithelial tight junctions during energetic stress via its effector GIV/Girdin.” Elife 5.

Biehler, C., L. T. Wang, M. Sevigny, A. Jette, C. L. Gamblin, R. Catterall, E. Houssin, L. McCaffrey and P. Laprise (2020). “Girdin is a component of the lateral polarity protein network restricting cell dissemination.” PLoS Genet 16(3): e1008674.

Boldt, K., J. van Reeuwijk, Q. Lu, K. Koutroumpas, T. M. Nguyen, Y. Texier, S. E. van Beersum, N. Horn, J. R. Willer, D. A. Mans, G. Dougherty, I. J. Lamers, K. L. Coene, H. H. Arts, M. J. Betts, T. Beyer, E. Bolat, C. J. Gloeckner, K. Haidari, L. Hetterschijt, D. Iaconis, D. Jenkins, F. Klose, B. Knapp, B. Latour, S. J. Letteboer, C. L. Marcelis, D. Mitic, M. Morleo, M. M. Oud, M. Riemersma, S. Rix, P. A. Terhal, G. Toedt, T. J. van Dam, E. de Vrieze, Y. Wissinger, K. M. Wu, G. Apic, P. L. Beales, O. E. Blacque, T. J. Gibson, M. A. Huynen, N. Katsanis, H. Kremer, H. Omran, E. van Wijk, U. Wolfrum, F. Kepes, E. E. Davis, B. Franco, R. H. Giles, M. Ueffing, R. B. Russell, R. Roepman and U. K. R. D. Group (2016). “An organelle-specific protein landscape identifies novel diseases and molecular mechanisms.” Nat Commun 7: 11491.

Breton, B., E. Sauvageau, J. Zhou, H. Bonin, C. Le Gouill and M. Bouvier (2010). “Multiplexing of multicolor bioluminescence resonance energy transfer.” Biophys J 99(12): 4037–4046.

Brown, N. E., N. A. Lambert and J. R. Hepler (2016). “RGS14 regulates the lifetime of Galpha-GTP signaling but does not prolong Gbetagamma signaling following receptor activation in live cells.” Pharmacol Res Perspect 4(5): e00249.

Choi, J. S., K. H. Kim, E. Oh, Y. K. Shin, J. Seo, S. H. Kim, S. Park and Y. L. Choi (2017). “Girdin protein expression is associated with poor prognosis in patients with invasive breast cancer.” Pathology 49(6): 618–626.

Coleman, B. D., A. Marivin, K. Parag-Sharma, V. DiGiacomo, S. Kim, J. S. Pepper, J. Casler, L. T. Nguyen, M. R. Koelle and M. Garcia-Marcos (2016). “Evolutionary Conservation of a GPCR-Independent Mechanism of Trimeric G Protein Activation.” Mol Biol Evol 33(3): 820–837.

de Opakua, A. I., K. Parag-Sharma, V. DiGiacomo, N. Merino, A. Leyme, A. Marivin, M. Villate, L. T. Nguyen, M. A. de la Cruz-Morcillo, J. B. Blanco-Canosa, S. Ramachandran, G. S. Baillie, R. A. Cerione, F. J. Blanco and M. Garcia-Marcos (2017). “Molecular mechanism of Galphai activation by non-GPCR proteins with a Galpha-Binding and Activating motif.” Nat Commun 8: 15163.

Ear, J., A. Saklecha, N. Rajapakse, J. Choi, M. Ghassemian, I. Kufareva and P. Ghosh (2020). “Tyrosine-Based Signals Regulate the Assembly of DaplePARD3 Complex at Cell-Cell Junctions.” iScience 23(2): 100859.

Enomoto, A., H. Murakami, N. Asai, N. Morone, T. Watanabe, K. Kawai, Y. Murakumo, J. Usukura, K. Kaibuchi and M. Takahashi (2005). “Akt/PKB regulates actin organization and cell motility via Girdin/APE.” Dev Cell 9(3): 389–402.

Ezan, J., L. Lasvaux, A. Gezer, A. Novakovic, H. May-Simera, E. Belotti, A. C. Lhoumeau, L. Birnbaumer, S. Beer-Hammer, J. P. Borg, A. Le Bivic, B. Nurnberg, N. Sans and M. Montcouquiol (2013). “Primary cilium migration depends on G-protein signalling control of subapical cytoskeleton.” Nat Cell Biol 15(9): 1107–1115.

Fanning, A. S. and J. M. Anderson (1999). “PDZ domains: fundamental building blocks in the organization of protein complexes at the plasma membrane.” J Clin Invest 103(6): 767–772.

Garcia-Marcos, M., P. Ghosh, J. Ear and M. G. Farquhar (2010). “A structural determinant that renders G alpha(i) sensitive to activation by GIV/girdin is required to promote cell migration.” J Biol Chem 285(17): 12765–12777.

Ghosh, P. (2015). “Heterotrimeric G proteins as emerging targets for network based therapy in cancer: End of a long futile campaign striking heads of a Hydra.” Aging (Albany NY) 7(7): 469–474.

Ghosh, P., A. O. Beas, S. J. Bornheimer, M. Garcia-Marcos, E. P. Forry, C. Johannson, J. Ear, B. H. Jung, B. Cabrera, J. M. Carethers and M. G. Farquhar (2010). “A G{alpha}i-GIV molecular complex binds epidermal growth factor receptor and determines whether cells migrate or proliferate.” Mol Biol Cell 21(13): 2338–2354.

Go, C. D., J. D. R. Knight, A. Rajasekharan, B. Rathod, G. G. Hesketh, K. T. Abe, J.-Y. Youn, P. Samavarchi-Tehrani, H. Zhang, L. Y. Zhu, E. Popiel, J.-P. Lambert, É. Coyaud, S. W. T. Cheung, D. Rajendran, C. J. Wong, H. Antonicka, L. Pelletier, B. Raught, A. F. Palazzo, E. A. Shoubridge and A.-C. Gingras (2019). “A proximity biotinylation map of a human cell.” bioRxiv: 796391.

Ha, A., A. Polyanovsky and T. Avidor-Reiss (2015). “Drosophila Hook-Related Protein (Girdin) Is Essential for Sensory Dendrite Formation.” Genetics 200(4): 1149–1159.

Houssin, E., U. Tepass and P. Laprise (2015). “Girdin-mediated interactions between cadherin and the actin cytoskeleton are required for epithelial morphogenesis in Drosophila.” Development 142(10): 1777–1784.

Ilyas, M., I. P. Tomlinson, A. Rowan, M. Pignatelli and W. F. Bodmer (1997). “Beta-catenin mutations in cell lines established from human colorectal cancers.” Proc Natl Acad Sci U S A 94(19): 10330–10334.

Jowett, T. (1999). “Analysis of protein and gene expression.” Methods Cell Biol 59: 63–85.

Kalogriopoulos, N. A., S. D. Rees, T. Ngo, N. J. Kopcho, A. V. Ilatovskiy, N. Sun, E. A. Komives, G. Chang, P. Ghosh and I. Kufareva (2019). “Structural basis for GPCR-independent activation of heterotrimeric Gi proteins.” Proc Natl Acad Sci U S A 116(33): 16394–16403.

Kannan, N. and V. W. Tang (2015). “Synaptopodin couples epithelial contractility to alpha-actinin-4-dependent junction maturation.” J Cell Biol 211(2): 407–434.

Karaman, R. and G. Halder (2018). “Cell Junctions in Hippo Signaling.” Cold Spring Harb Perspect Biol 10(5).

Ke, S., E. A. Alemu, C. Mertens, E. C. Gantman, J. J. Fak, A. Mele, B. Haripal, I. Zucker-Scharff, M. J. Moore, C. Y. Park, C. B. Vagbo, A. Kussnierczyk, A. Klungland, J. E. Darnell, Jr. and R. B. Darnell (2015). “A majority of m6A residues are in the last exons, allowing the potential for 3’ UTR regulation.” Genes Dev 29(19): 2037–2053.

Le-Niculescu, H., I. Niesman, T. Fischer, L. DeVries and M. G. Farquhar (2005). “Identification and characterization of GIV, a novel Galpha i/s-interacting protein found on COPI, endoplasmic reticulum-Golgi transport vesicles.” J Biol Chem 280(23): 22012–22020.

Liu, H., M. A. Flores, J. Meng, L. Zhang, X. Zhao, M. K. Rao, Y. Chen and Y. Huang (2015). “MeT-DB: a database of transcriptome methylation in mammalian cells.” Nucleic Acids Res 43(Database issue): D197–203.

Liu, H., H. Wang, Z. Wei, S. Zhang, G. Hua, S. W. Zhang, L. Zhang, S. J. Gao, J. Meng, X. Chen and Y. Huang (2018). “MeT-DB V2.0: elucidating context-specific functions of N6-methyl-adenosine methyltranscriptome.” Nucleic Acids Res 46(D1): D281–D287.

Manjunath, G. P., P. L. Ramanujam and S. Galande (2018). “Structure function relations in PDZ-domain-containing proteins: Implications for protein networks in cellular signalling.” J Biosci 43(1): 155–171.

Marivin, A. and M. Garcia-Marcos (2019). “DAPLE and MPDZ bind to each other and cooperate to promote apical cell constriction.” Mol Biol Cell 30(16): 1900–1910.

McHenry, M. J. and S. M. van Netten (2007). “The flexural stiffness of superficial neuromasts in the zebrafish (Danio rerio) lateral line.” J Exp Biol 210(Pt 23): 4244–4253.

Metcalfe, W. K., C. B. Kimmel and E. Schabtach (1985). “Anatomy of the posterior lateral line system in young larvae of the zebrafish.” J Comp Neurol 233(3): 377–389.

Midde, K. K., N. Aznar, M. B. Laederich, G. S. Ma, M. T. Kunkel, A. C. Newton and P. Ghosh (2015). “Multimodular biosensors reveal a novel platform for activation of G proteins by growth factor receptors.” Proc Natl Acad Sci U S A 112(9): E937–946.

Mykytyn, K. and C. Askwith (2017). “G-Protein-Coupled Receptor Signaling in Cilia.” Cold Spring Harb Perspect Biol 9(9).

Nachury, M. V. and D. U. Mick (2019). “Establishing and regulating the composition of cilia for signal transduction.” Nat Rev Mol Cell Biol 20(7): 389–405.

Nechipurenko, I. V., A. Olivier-Mason, A. Kazatskaya, J. Kennedy, I. G. McLachlan, M. G. Heiman, O. E. Blacque and P. Sengupta (2016). “A Conserved Role for Girdin in Basal Body Positioning and Ciliogenesis.” Dev Cell 38(5): 493–506.

Ohara, K., A. Enomoto, T. Kato, T. Hashimoto, M. Isotani-Sakakibara, N. Asai, M. Ishida-Takagishi, L. Weng, M. Nakayama, T. Watanabe, K. Kato, K. Kaibuchi, Y. Murakumo, Y. Hirooka, H. Goto and M. Takahashi (2012). “Involvement of Girdin in the determination of cell polarity during cell migration.” PLoS One 7(5): e36681.

Oshita, A., S. Kishida, H. Kobayashi, T. Michiue, T. Asahara, M. Asashima and A. Kikuchi (2003). “Identification and characterization of a novel Dvl-binding protein that suppresses Wnt signalling pathway.” Genes Cells 8(12): 1005–1017.

Pan, C. Q., M. Sudol, M. Sheetz and B. C. Low (2012). “Modularity and functional plasticity of scaffold proteins as p(l)acemakers in cell signaling.” Cell Signal 24(11): 2143–2165.

Park, T. J., B. J. Mitchell, P. B. Abitua, C. Kintner and J. B. Wallingford (2008). “Dishevelled controls apical docking and planar polarization of basal bodies in ciliated epithelial cells.” Nat Genet 40(7): 871–879.

Pawson, T. and P. Nash (2003). “Assembly of cell regulatory systems through protein interaction domains.” Science 300(5618): 445–452.

Rausch, V. and C. G. Hansen (2020). “The Hippo Pathway, YAP/TAZ, and the Plasma Membrane.” Trends Cell Biol 30(1): 32–48.

Rouaud, F., S. Sluysmans, A. Flinois, J. Shah, E. Vasileva and S. Citi (2020). “Scaffolding proteins of vertebrate apical junctions: structure, functions and biophysics.” Biochim Biophys Acta Biomembr 1862(10): 183399.

Sasaki, K., T. Kakuwa, K. Akimoto, H. Koga and S. Ohno (2015). “Regulation of epithelial cell polarity by PAR-3 depends on Girdin transcription and Girdin-Galphai3 signaling.” J Cell Sci 128(13): 2244–2258.

Scott, J. D. and T. Pawson (2009). “Cell signaling in space and time: where proteins come together and when they’re apart.” Science 326(5957): 1220–1224.

Shevchenko, A., M. Wilm, O. Vorm and M. Mann (1996). “Mass spectrometric sequencing of proteins silver-stained polyacrylamide gels.” Anal Chem 68(5): 850–858.

Siletti, K., B. Tarchini and A. J. Hudspeth (2017). “Daple coordinates organ-wide and cell-intrinsic polarity to pattern inner-ear hair bundles.” Proc Natl Acad Sci U S A 114(52): E11170–E11179.

Subbaiah, V. K., C. Kranjec, M. Thomas and L. Banks (2011). “PDZ domains: the building blocks regulating tumorigenesis.” Biochem J 439(2): 195–205.

Tang, V. W. and W. M. Brieher (2013). “FSGS3/CD2AP is a barbed-end capping protein that stabilizes actin and strengthens adherens junctions.” J Cell Biol 203(5): 815–833.

Teo, G., G. Liu, J. Zhang, A. I. Nesvizhskii, A. C. Gingras and H. Choi (2014). “SAINTexpress: improvements and additional features in Significance Analysis of INTeractome software.” J Proteomics 100: 37–43.

Vermeulen, S. J., E. A. Bruyneel, M. E. Bracke, G. K. De Bruyne, K. M. Vennekens, K. L. Vleminckx, G. J. Berx, F. M. van Roy and M. M. Mareel (1995). “Transition from the noninvasive to the invasive phenotype and loss of alpha-catenin in human colon cancer cells.” Cancer Res 55(20): 4722–4728.

Volff, J. N. (2005). “Genome evolution and biodiversity in teleost fish.” Heredity (Edinb) 94(3): 280–294.

Wang, S., Y. Lei, Z. Cai, X. Ye, L. Li, X. Luo and C. Yu (2018). “Girdin regulates the proliferation and apoptosis of pancreatic cancer cells via the PI3K/Akt signalling pathway.” Oncol Rep 40(2): 599–608.

Wang, W., H. Chen, W. Gao, S. Wang, K. Wu, C. Lu, X. Luo, L. Li and C. Yu (2020). “Girdin interaction with vimentin induces EMT and promotes the growth and metastasis of pancreatic ductal adenocarcinoma.” Oncol Rep 44(2): 637–649.

Wang, Y., J. V. Denisova, K. S. Kang, J. D. Fontes, B. T. Zhu and A. B. Belousov (2010). “Neuronal gap junctions are required for NMDA receptor-mediated excitotoxicity: implications in ischemic stroke.” J Neurophysiol 104(6): 3551–3556.

Wong, V. (1997). “Phosphorylation of occludin correlates with occludin localization and function at the tight junction.” Am J Physiol 273(6): C1859–1867.

Yang, Z., F. Yang, Y. Zhang, X. Wang, J. Shi, H. Wei, F. Sun and Y. Yu (2018). “Girdin protein: A potential metastasis predictor associated with prognosis in lung cancer.” Exp Ther Med 15(3): 2837–2843.

Zaccara, S., R. J. Ries and S. R. Jaffrey (2019). “Reading, writing and erasing mRNA methylation.” Nat Rev Mol Cell Biol 20(10): 608–624.

Zhu, L. Y., Y. R. Zhu, D. J. Dai, X. Wang and H. C. Jin (2018). “Epigenetic regulation of alternative splicing.” Am J Cancer Res 8(12): 2346–2358.

