## Supplemental File for "Evolution of Modularity, Interactome and Functions of GIV/Girdin (CCDC88A) from Invertebrates to Vertebrates"

#### SUPPLEMENTARY FIGURE LEGENDS

##### Figure S1 – Supplement to Figure 1
**The C-terminus of GIV has an evolutionarily conserved functional PDZ-binding motif downstream of its G protein binding and/or modulatory domains.**

**A)** Table summarizing the characterized modules and motifs in GIV and Daple. "P" indicates presence, "X" indicates no description, "-" indicates absence.

**B)** Amino acid alignment of the PBM across various species.

**C)** A magnified image of a zebrafish stained for zGIV (CCDC88Ab; from Figure 1C) is shown.

##### Figure S2- Supplement to Figure 4

**The PBM motif in GIV-L is functional and binds PDZ-proteins ParD3 and Dvl.**

**A-C)** GST-pulldown assays were carried out using purified GST-tagged PDZ domains of ParD3 (A) or Dvl (B). Bound proteins were visualized (left) and equal loading of cell lysates (right) were confirmed by immunoblotting (IB). Lysates of HEK293T cells (C) exogenously expressing myc-tagged GIV (wt or F1685A) or GIV-L (wt, F1685A, ΔPBM, or F1685A/ΔPBM double mutant) that were used as source of GIV proteins for the pulldown assays.

**Figure S3 – Supplement to Figure 6**

**GIV-L's 'PDZ-ome' provides clues into specific junction-sensing pathways GIV-L may modulate.**

Reactome pathway analysis was performed on the GIV-L bound 'PDZ-ome' (**Fig 6D'-E'**) and findings are visualized as ReacFoam (top) or table of statistically enriched pathways (bottom). Boxed regions on top are magnified. Arrows (red) highlight the overrepresentation of two specific junction-sensing pathways, NMDA and HIPPO.

**Figure S4 – Supplement to Figure 8**

**Validation studies for customized rabbit polyclonal antibodies used to detect GIV and GIV-L individually.**

**A)** Schematic depicts the ectopically expressed GIV or GIV-L construct in HEK293T cells and the binging region of the antibodies used.

**B)** Various GIV antibodies was used to immunoblot HEK293T cell lysates overexpressing EGFP-tagged GIV or GIV-L. EGFP-tagged GIV-L (CT) was overexpressed in HEK293T cells and purified using an anti-GFP camelid antibody. Purified protein, along with cell lysates, was used in SDS-PAGE and western blotting analysis to validate specificity of GIV antibodies.

SUPPLEMENTARY FIGURES

### Figure S1

**A**

| | HOOK | | Coiled-coil | | GBD | | GEM | | PBM | |
|---|---|---|---|---|---|---|---|---|---|---|
| **Drosophila** | ✓ | - | ✓ | - | X | - | X | - | ✓ | - |
| **C. elegans** | ✓ | - | ✓ | - | ✓ | - | X | - | ✓ | - |
| **Zebrafish** | ✓ | ✓ | ✓ | ✓ | ✓ | ✓ | ✓ | ✓ | ✓ | ✓ |
| **Lizard** | ✓ | ✓ | ✓ | ✓ | ✓ | ✓ | ✓ | ✓ | ✓ | ✓ |
| **Chicken** | ✓ | ✓ | ✓ | ✓ | ✓ | ✓ | ✓ | ✓ | ✓ | ✓ |
| **Dog** | ✓ | ✓ | ✓ | ✓ | ✓ | ✓ | ✓ | ✓ | X | ✓ |
| **Mouse** | ✓ | ✓ | ✓ | ✓ | ✓ | ✓ | ✓ | ✓ | X | ✓ |
| **Monkey** | ✓ | ✓ | ✓ | ✓ | ✓ | ✓ | ✓ | ✓ | X | ✓ |
| **Human** | ✓ | ✓ | ✓ | ✓ | ✓ | ✓ | ✓ | ✓ | X | ✓ |
| | GIV | Daple | GIV | Daple | GIV | Daple | GIV | Daple | GIV | Daple |

**B**

| | |
|---|---|
| *Homo sapiens* | QTVWYEYGCI |
| *Pan troglodytes* | QTVWYEYGCI |
| *Rattus norvegicus* | QTVWYEYGCI |
| *Mus musculus* | QIVWYEYGCI |
| *Gallus gallus* | KTVWYEYGCV |
| *Pogona vitticeps* | QTIWYEYGCV |
| *Xenopus laevis* | QSIWYEYGCV |
| *Danio rerio* | DGLWYEYGCV |
| *Drosophila melanogaster* | NSIWYEYGCV |
| *Caenorhabditis elegans* | STIWHEYGCV |
| | ...*.****. |

**C**

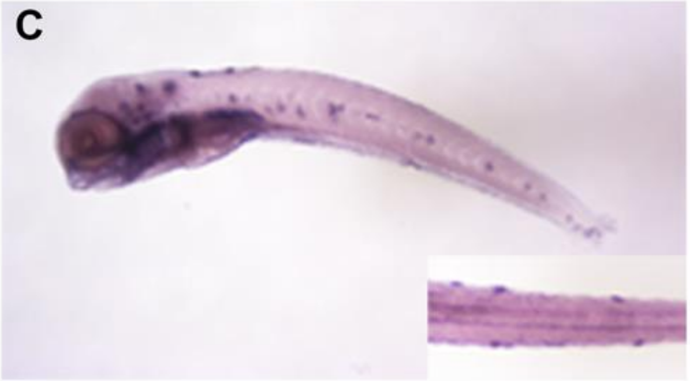

### Figure S2

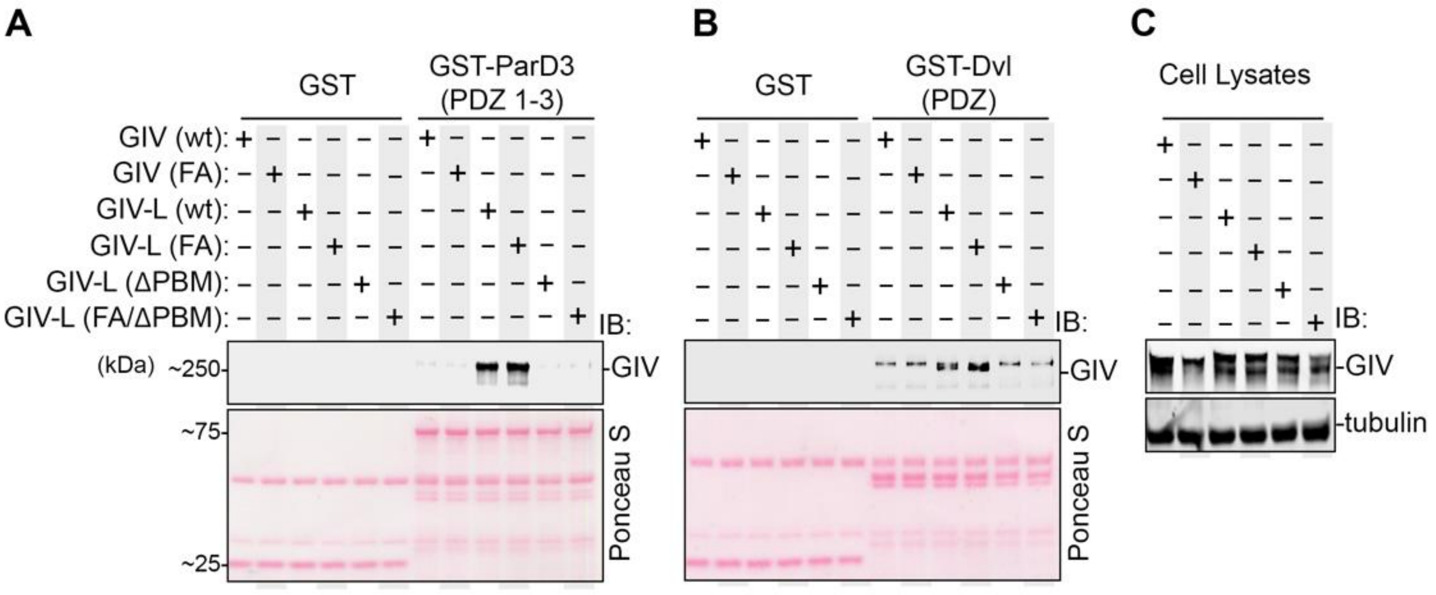
A
GST
GST-ParD3 (PDZ 1-3)
GIV (wt):
GIV (FA):
GIV-L (wt):
GIV-L (FA):
GIV-L (ΔPBM):
GIV-L (FA/ΔPBM):
IB:
(kDa)
~250
GIV
~75
~25
Ponceau S
B
GST
GST-Dvl (PDZ)
IB:
GIV
Ponceau S
C
Cell Lysates
IB:
GIV
tubulin

### Figure S3

Reacfome analysis of GIV-L's "PDZ-ome"

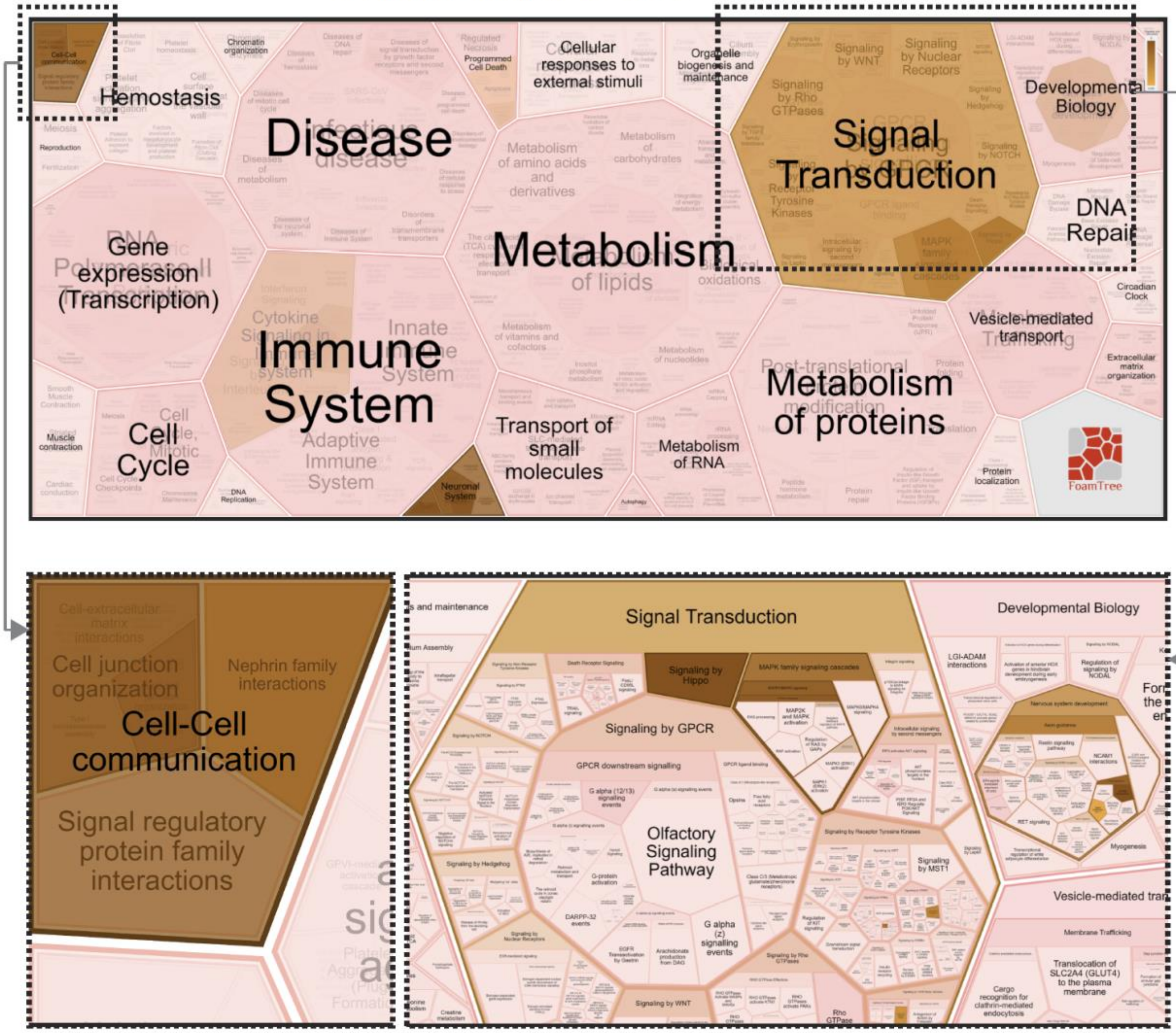


| Pathway name | Entities found | Entities Total | Entities ratio | Entities pValue | Entities FDR | Reactions found | Reactions total | Reactions ratio | Species name |
|---|---|---|---|---|---|---|---|---|---|
| Assembly and cell surface presentation of NMDA receptors | 5 | 49 | 0.003 | 3.15E-8 | 4.54E-6 | 7 | 23 | 0.002 | Homo sapiens |
| Ras activation upon Ca2+ influx through NMDA receptor | 4 | 27 | 0.002 | 1.95E-7 | 8.09E-6 | 2 | 2 | 0 | Homo sapiens |
| Negative regulation of NMDA receptor-mediated neuronal transmission | 4 | 27 | 0.002 | 1.95E-7 | 8.09E-6 | 1 | 4 | 0 | Homo sapiens |
| Unblocking of NMDA receptors, glutamate binding and activation | 4 | 28 | 0.002 | 2.25E-7 | 8.09E-6 | 5 | 5 | 0 | Homo sapiens |
| Long-term potentiation | 4 | 31 | 0.002 | 3.36E-7 | 1E-5 | 3 | 7 | 0.001 | Homo sapiens |
| CREB1 phosphorylation through NMDA receptor-mediated activation of RAS signaling | 4 | 39 | 0.003 | 8.34E-7 | 2E-5 | 2 | 7 | 0.001 | Homo sapiens |
| Activation of NMDA receptors and postsynaptic events | 5 | 114 | 0.008 | 1.98E-6 | 4E-5 | 18 | 71 | 0.005 | Homo sapiens |
| Protein-protein interactions at synapses | 4 | 93 | 0.006 | 2.52E-5 | 4.54E-4 | 13 | 33 | 0.003 | Homo sapiens |
| Post NMDA receptor activation events | 4 | 96 | 0.007 | 2.85E-5 | 4.56E-4 | 5 | 39 | 0.003 | Homo sapiens |
| Neurotransmitter receptors and postsynaptic signal transmission | 5 | 232 | 0.016 | 6E-5 | 8.35E-4 | 21 | 109 | 0.008 | Homo sapiens |
| NrCAM interactions | 2 | 7 | 0 | 8E-5 | 1.03E-3 | 1 | 4 | 0 | Homo sapiens |
| RUNX1 regulates expression of components of tight junctions | 2 | 8 | 0.001 | 1.03E-4 | 1.24E-3 | 2 | 6 | 0 | Homo sapiens |
| Neurexins and neuroligins | 3 | 60 | 0.004 | 1.86E-4 | 1.95E-3 | 12 | 19 | 0.001 | Homo sapiens |
| Apoptotic cleavage of cell adhesion proteins | 2 | 11 | 0.001 | 1.95E-4 | 1.95E-3 | 2 | 10 | 0.001 | Homo sapiens |
| Activation of Ca-permeable Kainate Receptor | 2 | 13 | 0.001 | 2.71E-4 | 2.44E-3 | 2 | 2 | 0 | Homo sapiens |
| Ionotropic activity of kainate receptors | 2 | 14 | 0.001 | 3.15E-4 | 2.83E-3 | 2 | 4 | 0 | Homo sapiens |
| Transmission across Chemical Synapses | 5 | 352 | 0.024 | 4.13E-4 | 3.3E-3 | 23 | 162 | 0.012 | Homo sapiens |
| Signaling by Hippo | 2 | 22 | 0.001 | 7.7E-4 | 5.88E-3 | 4 | 30 | 0.002 | Homo sapiens |
| Synaptic adhesion-like molecules | 2 | 23 | 0.002 | 8.4E-4 | 5.88E-3 | 1 | 8 | 0.001 | Homo sapiens |
| L1CAM interactions | 3 | 130 | 0.009 | 1.73E-3 | 1.09E-2 | 2 | 54 | 0.004 | Homo sapiens |

#### Figure S4

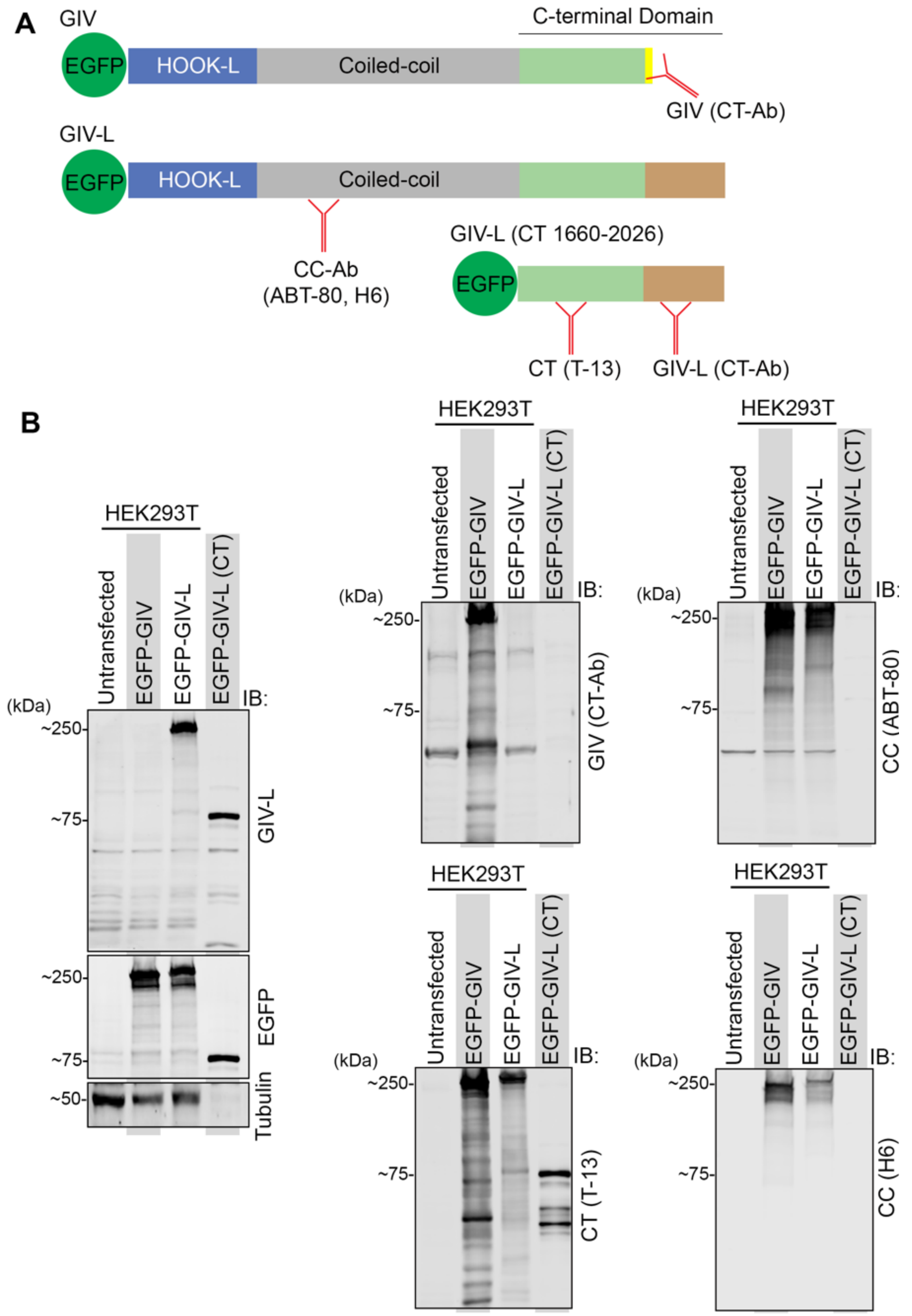

A
GIV
C-terminal Domain
EGFP
HOOK-L
Coiled-coil
GIV (CT-Ab)
GIV-L
EGFP
HOOK-L
Coiled-coil
CC-Ab
(ABT-80, H6)
GIV-L (CT 1660-2026)
EGFP
CT (T-13)
GIV-L (CT-Ab)
B
HEK293T
Untransfected
EGFP-GIV
EGFP-GIV-L
EGFP-GIV-L (CT)
IB:
(kDa)
~250-
~75-
~50-
GIV-L
EGFP
Tubulin
GIV (CT-Ab)
CC (ABT-80)
CT (T-13)
CC (H6)
